# A branching model of cell fate decisions in the enteric nervous system

**DOI:** 10.1101/2022.07.12.499640

**Authors:** Anna Laddach, Song Hui Chng, Reena Lasrado, Fränze Progatzky, Michael Shapiro, Artem Artemov, Marisol Sampedro Castaneda, Alek Erickson, Ana Carina Bon-Frauches, Jens Kleinjung, Stefan Boeing, Sila Ultanir, Igor Adameyko, Vassilis Pachnis

## Abstract

How neurogenesis and gliogenesis are coordinated during development and why mature glial cells often share properties with neuroectodermal progenitors remains unclear. Here, we have used single cell RNA sequencing to map the regulatory landscape of neuronal and glial differentiation in the mammalian enteric nervous system (ENS). Our analysis indicates that neurogenic trajectories branch directly from a linear gliogenic axis defined by autonomic neural crest cells adopting sequential states as they progressively lose their strong neurogenic bias and acquire properties of adult enteric glia. We identify gene modules associated with transcriptional programs driving enteric neurogenesis and cell state transitions along the gliogenic axis. By comparing the chromatin accessibility profile of autonomic neural crest and adult enteric glia we provide evidence that the latter maintain an epigenetic memory of their neurogenic past. Finally, we demonstrate that adult enteric glia maintain neurogenic potential and are capable of generating enteric neurons in certain contexts by activating transcriptional programs employed by early ENS progenitors. Our studies uncover a novel configuration of enteric neurogenesis and gliogenesis that enables the coordinate development of ENS lineages and provides a mechanistic explanation for the ability of enteric glia to be functionally integrated into the adult intestine and simultaneously maintain attributes of early ENS progenitors.

## Introduction

Neural crest cells are multipotential embryonic progenitors that generate diverse tissues in vertebrates, including the autonomic and somatosensory ganglia of the peripheral nervous system (PNS) and musculoskeletal structures^1^. Intense investigation over several decades has identified molecular and signalling pathways that regulate the development of neural crest-derived tissues^2^. More recently, transcriptional profiling at single cell resolution has provided evidence that major neural crest lineages (autonomic, sensory, mesenchymal) emerge as undifferentiated progenitors make binary cell fate choices by resolving competing transcriptional programmes^3^. Although neurogenesis and gliogenesis in the PNS are also formalised as alternative and irreversible cell fate decisions made by common progenitors, the strategies employed for the timely and balanced generation of peripheral neurons and glia, remain unclear.

The enteric nervous system (ENS), is the branch of the autonomic nervous system that encompasses all neurons and glial cells intrinsic to the gut wall. Diverse classes of enteric neurons and glia are organised into highly complex circuits that are distributed throughout the length of the gut and control, among other functions, intestinal motility, epithelial secretions and blood flow^4^. By responding to cues from the microbiota and establishing functional interactions with epithelial and immune cells in the gut wall, the ENS plays key roles in regulating intestinal immunity and host defence^5–7^. Extending these studies, our recent work has shown that interferon-γ (IFNγ) signalling in enteric glial cells (EGCs) regulates the immune response and tissue repair following helminth infection of the intestine^8^. Interestingly, the IFNγ-enteric glia signalling axis is also critical for maintaining immune homeostasis in the mammalian gut at steady state^8^, suggesting that the immune potential of enteric glia is integral to the differentiation and maturation of this lineage.

The vast majority of adult ENS cells are derived from a small population of autonomic neural crest cells (ANCCs) which invade the foregut during embryogenesis. Earlier studies, including our own lineage tracing experiments and clonal analysis, demonstrated that the Sox10-expressing descendants of the founder neural crest cell population represent the bipotential self-renewing progenitors that fuel the expansion of the ENS during development and give rise to all types of enteric neurons and glia^9–11^. Transcriptional profiling of the ENS during development has provided evidence that enteric neuron subtype diversity is built on the cardinal specification of postmitotic neuronal precursors into nitrergic and cholinergic subtypes, which are further diversified in response to subtype-specific transcriptional regulators^12–16^. Nevertheless, it remains unclear whether the molecular mechanisms promoting cell cycle exit and neuronal commitment of undifferentiated ENS progenitors also specify neuronal subtype identity. Although glial differentiation is thought to be initiated at relatively early stages of ENS development, identifying key milestones of enteric gliogenesis and the associated intermediate cell types, has been problematic given that the majority of molecular markers used for the identification of adult enteric glia are also expressed by ANCCs and their undifferentiated progeny in the gut. Consequently, how the neurogenic and gliogenic output of undifferentiated ENS progenitors is regulated and integrated across developmental time remains unclear. Intriguingly, despite being considered a terminally differentiated cell type fully integrated into the tissue circuitry of the adult intestine, vertebrate EGCs are capable of differentiating into neurons, either constitutively (zebrafish) or under special experimental conditions (rodents)^10, 17, 18^, suggesting that they preserve the neurogenic potential of their progenitors. Nevertheless, it remains unclear whether neurogenesis by adult EGCs is controlled by the neurogenic programmes that normally drive the differentiation of Sox10^+^ ENS progenitors into neurons during development.

To understand how neuronal and glial cell fate decisions are implemented during PNS development, we characterised the transcriptional and chromatin accessibility profile of Sox10-expressing cells during ENS development and adulthood. We demonstrate that the founder population of neural crest cells and early ENS progenitors are characterised by a strong neurogenic bias reflected in their collective transcriptional and epigenetic profile. As development proceeds, similar programmes of neuronal differentiation are activated, albeit with diminishing efficiency, by ENS progenitors as they reduce their neurogenic output and simultaneously acquire properties associated with mature enteric glia. Despite the lack of neurogenesis in the adult ENS, enteric glia maintain an epigenetic memory of their neurogenic past and are capable of conditionally reactivating gene expression programmes that are normally employed to drive neuronal differentiation during development. Our studies advance the understanding of how neurogenesis and gliogenesis are coordinated across ENS development and uncover mechanisms that could be harnessed for restoring neural deficiencies in the peripheral and central nervous system.

## Results

### Transcriptional profile of ENS cells during development

To investigate the establishment of the mammalian ENS lineages, we performed an integrated analysis of novel and previously generated^8, 11^ scRNA-seq datasets of tdTomato (tdT) labelled Sox10^+^ cells isolated from the small intestine of Sox10CreER|tdT reporter mice^8, 10^ during development and adulthood (Fig. 1a; see Materials and Methods). Uniform manifold approximation and projection (UMAP) analysis distributed cells according to developmental stage (Fig. 1b; Extended Data Fig. 1a). tdT^+^ cells from E13.5 embryos formed a proliferative cluster co-expressing undifferentiated ENS progenitor (*Erbb3*, *Sox10*, *Fabp7* and *Plp1*) and neuronal (*Tubb3*, *Elavl4*, *Ret* and *Phox2b*) markers (called hereafter early ENS progenitors-eEPs), and an adjoining and largely post-mitotic cluster of committed neuronal precursors, which downregulated the progenitor and upregulated neuronal markers (early enteric neuronal precursors-eENPs) (Fig. 1b, c; Extended Data Fig. 1b). E17.5 and P1 cells, were clearly separated from their E13.5 counterparts and formed adjoining clusters of ENS progenitors (late ENS progenitors 1 and 2-lEP1 and lEP2) and a common cluster of committed neuronal precursors (late enteric neuronal precursors-lENPs) which bridged lEPs with adult enteric neurons sequenced in parallel (Fig. 1b). Relative to eEPs, lEPs were characterised by reduced proliferative activity, downregulation of neuronal genes and uniform upregulation of the glial markers *S100b* and *Plp1* (Fig. 1c; Extended Data Fig. 1b, c), indicating that ENS progenitors from late prenatal stages display a glial-like character and reduced neurogenic output. This was supported by k-means clustering (k=2) with the progenitor and neurogenic gene markers, which showed that the representation of committed neural precursors in the tdT^+^ cell population was progressively reduced from E13.5 to P1 (Fig. 1d). Finally, tdT^+^ cells from P26 and P61 animals were intermixed in two quiescent clusters of glial cells (EGC1 and EGC2)^8^ with no committed neuronal precursors emerging from them (Fig. 1b, c; Extended Data Fig. 1b), consistent with the lack of neurogenesis in adult mammalian ENS under steady state conditions^10, 19^. Therefore, during development, the cellular landscape of the mammalian ENS is shaped by time-dependent changes in the proliferative activity, transcriptional profile and neurogenic output of Sox10^+^ progenitors.

**Figure 1.**
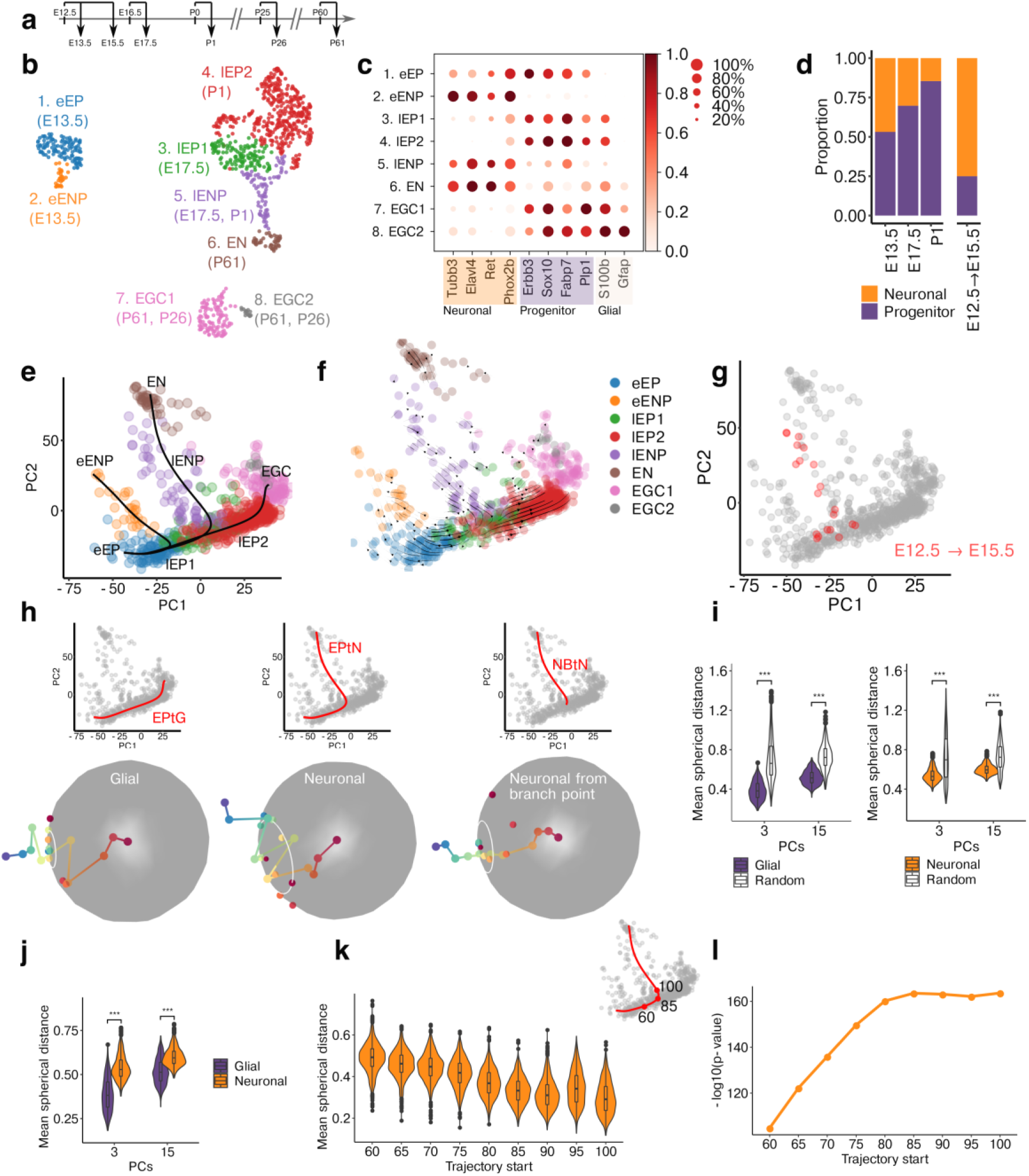
scRNA-seq and TrajectoryGeometry support a branching model of ENS lineage development. **a,** Developmental timeline for cell labelling and isolation of SOX10^+^ ENS cells. **b,** UMAP representation of sequenced cells coloured by cluster. **c,** Dot plot representing the level of expression of neuronal, progenitor and glial markers in the clusters shown in **b**. The colour scale represents the mean expression level, and dot size represents the percentage of cells with non-zero expression within a given cluster. **d,** Stacked bar plot showing the percentage of neuronal and progenitor fractions within the cell populations isolated at the indicated timepoints. **e,** Slingshot analysis indicating the differentiation trajectories of ENS lineages. eEP to eENP, early neurogenic trajectory; eEP to EN, late neurogenic trajectory; eEP to EGC, gliogenic trajectory. **f,** PCA plot shown with velocity field suggesting the direction of cell differentiation. **g,** PCA plot indicating tdT^+^ ENS cells labelled at E12.5 and sequenced at E15.5. Lineage traced cells (red) are intermixed with lEPs and lEPNs (grey). **h,** Individual paths for the eEP to Glia (EPtG), eEP to Neuron (EPtN) and late Neuronal branch to Neuron (NbtN) trajectories shown on the PCA plot (top) and projected onto a sphere (bottom). The radius of the white circles indicates the mean spherical distance from the centre of the projections. **i,** Violin plots indicating the mean spherical distance (radii of the white circles in **h**) for paths sampled from the EPtG and EPtN trajectories (purple and orange, respectively) relative to random trajectories (white). Statistics calculated using 1000 random paths from each trajectory. **j,** Violin plots indicating the mean spherical distance of the EPtG (purple) and EPtN (orange) trajectories. **k,** Violin plots indicating the mean spherical distance for the EPtN trajectory starting from successively later points in pseudotime, as the branch point is approached (85 value on the EPtN trajectory shown in the top right inset). **l,** Line graph indicating the –log10(p-value) for the significance of directionality for the neuronal trajectory relative to random trajectories, starting from successively later points in pseudotime.

### Branching model of cell fate decisions in the ENS

To delineate the cell differentiation trajectories of the ENS, we analysed our transcriptomic datasets using lineage and pseudotime inference algorithms^20, 21^. Consistent with our previous clonal analysis^11^, we identified a time-axis aligned eEP to glia (EPtG) trajectory and eEP to neuron (EPtN) trajectories emerging orthogonally from it (Fig. 1e, f). To provide experimental support for the inferred trajectories, we performed scRNA-seq on E15.5 ENS cells from Sox10CreER|tdT embryos that were labelled with tdT 3 days earlier (Fig. 1a). Despite labelling at E12.5, all lineage traced cells clustered with lEP1 and lENP (Fig. 1g; Extended Data Fig. 1d), were composed mainly of committed neuronal precursors (Fig. 1d) and showed reduced proliferative activity (Extended Data Fig. 1b). These findings support the strong neurogenic bias of eEPs and suggest that during development a common pool of ENS progenitors and derivative neuronal precursors transition collectively to distinct and sequentially emerging transcriptional states.

Based on the configuration of ENS differentiation trajectories, we postulated that the neurogenic axes branch off a gliogenic backbone that maintains a relatively consistent directionality of gene expression change throughout development. To quantify this directionality, we developed an R package (TrajectoryGeometry; https://bioconductor.org/packages/devel/bioc/html/TrajectoryGeometry.html), which allowed us to sample pseudotime paths along an arbitrary number of dimensions of the principal component analysis space and project them onto an n-dimensional sphere^22^. We reasoned that the more consistent the directionality of a path, the smaller the radius of the circle representing the mean spherical distance of projected points from a reference centre (Extended Data Fig. 2a; see also Materials and Methods). Sampling 1000 paths over the same developmental time (E13.5 to Adult) revealed significant directionality of both the EPtG and EPtN trajectories relative to an equal number of randomized paths (Fig. 1h, i; Extended Data Fig. 2b; Wilcoxon test, p < 0.001). However, direct comparison of these trajectories indicated that the EPtG axis maintained a more consistent directionality of gene expression change relative to EPtN (Fig. 1h, j, Extended Data Fig. 2b; U-test, p < 0.001). Furthermore, the initial decrease and subsequent stabilisation of the mean distance values for EPtN segments commencing at successively later points in pseudotime supported the presence of a branch point (corresponding to the inflection point at pseudotime value of ∼85), after which the directionality of the trajectory becomes more significant (Fig. 1h, k, l, Extended Data Fig. 2c). A similar pattern was not evident for the EPtG axis (Extended Data Fig. 2c), indicating that this trajectory does not change course. As expected, genes positively associated with the directionality of EPtG and EPtN included glial markers (*Apoe*, *S100b*) and neuronal markers (*Elavl3*, *Tubb3*), while negatively associated genes included several cell cycle regulators (*Top2a*, *Mki67*) (Extended Data Fig. 2d), indicating that cell proliferation is reduced along both differentiation trajectories. Together, our experimental and computational analyses suggest that, unlike the binary choices made by early neural crest cells, ENS progenitors either exit the gliogenic trajectory by committing to a neurogenic path or remain on course giving rise by default to the mature enteric glial cell state.

### Common molecular mechanisms underlie enteric neurogenesis during development

The egress of neurogenic paths from advancing positions along the EPtG axis (Fig. 1e) led us to wonder about the relationship of the transcriptional mechanisms driving enteric neuron commitment and differentiation at sequential branching points. To explore this, we used ANTLER (Another Transcriptome Lineage Explorer)^23^ (https://juliendelile.github.io/Antler/) to identify sets of genes with co-ordinate patterns of expression across the data (Extended Data Fig. 3; Suppl. Table 1). Focusing on gene modules (GMs) expressed by committed neuronal precursors, we noted that all GMs expressed by lENPs were also expressed by eENPs (Fig. 2a; Extended Data Fig. 3). These included GMs related to neurodevelopmental processes (GMs 26, 23, 8 and 47) and cardinal subtype specification of enteric neurons (GMs 65 and 55, corresponding to nitrergic and cholinergic neurons, respectively), although the sparser expression of these neuronal modules in eENPs suggested a less mature phenotype (Fig. 2a, b). The more mature character of lENPs was also supported by the upregulation of GM48, which includes gene markers (such as *Calb2* and *Tac1*) for neuronal subtypes emerging at advanced positions along the cholinergic branch of enteric neuron specification^16^ (Fig. 2a, b). To further explore the transcriptional mechanisms driving enteric neurogenesis at different developmental stages, we also analysed the expression of transcription factors identified using Monocle’s gene test function (differentialGeneTest)^24, 25^. In general, these transcription factors, which included known regulators of enteric neuron differentiation (*Sox10*, *Foxd3*, *Hmga2*, *Phox2a*, *Pbx3*), exhibited similar profiles between early and late neurogenesis (Fig. 2c). Finally, GM11, which included genes encoding Notch signalling ligands (*Dll1*, *Dll3*) and effectors (*Hes6*) promoting neurogenesis^26, 27^, was upregulated in committed enteric neuronal precursors emerging at different stages of mouse and human development (Fig. 2a; Extended Data Fig. 4, Extended Data Table 1 and 2), suggesting a ubiquitous role of Notch signalling in neuronal commitment and subtype specification of ENS progenitors.

**Figure 2.**
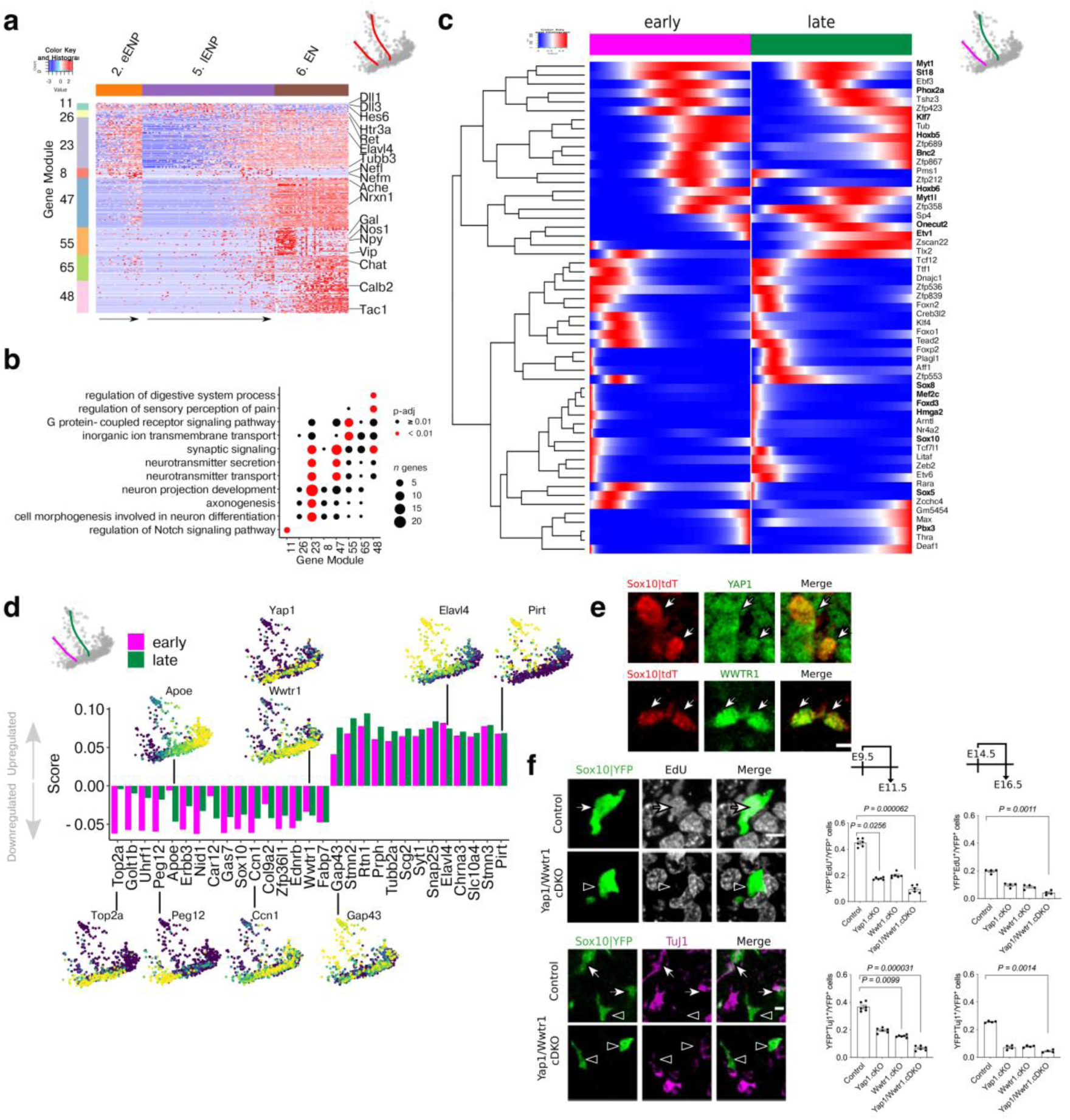
Transcriptomic analysis of early and late neurogenic trajectories of the ENS. **a,** Heatmap (scaled normalised expression) for selected gene modules (left) associated with eENPs, lENPs and ENs (top). Representative genes are indicated on the right. Inset indicates the early and late neurogenic trajectories on the PCA plot after the branch points. **b,** Dot plot showing statistical significance (colour of dot) and size of overlap (size of dot) between selected gene modules and indicated GO terms. **c,** Smoothed expression profiles for genes differentially expressed over pseudotime for early (magenta) or late (green) neurogenic trajectories from the branch points onwards. **d,** Bar plot showing the union of the top 10 TrajectoryGeometry genes positively associated (positive score) and top 10 genes negatively associated (negative score) with the directionality of early (magenta) and late (green) neurogenic trajectories. The PCA plots of the scRNA-seq dataset indicate the expression of selected genes (yellow denotes high expression). **e,** Immunostaining of tdT^+^ ENS cells (red; from E11.5 Sox10CreER|tdT mice) for YAP1 (top, green) and WWTR1 (bottom, green). Arrows show ENS cells that express indicated markers. **f,** Quantification of proliferation (EdU incorporation) and neuronal differentiation (TuJ1 immunostaining) of SOX10^+^ cells from control or mutant Sox10CreER|YFP mice carrying single conditional deletions of *Yap1* (Yap1.cKO) or *Wwtr1* (Wwtr1.cKO) or combined conditional deletions of *Yap1* and *Wwtr1* (Yap1/Wwtr1.cDKO). Deletions were induced either at E9.5 or E14.5 and guts analysed at E11.5 or E16.5, respectively. YFP^+^ cells and early neurons were identified by immunostaining for GFP (green) and TuJ1 (red), respectively. Incorporation of EdU is shown by the white dots. Arrows show ENS cells with EdU/TUJ1 signal, arrowheads show ENS cells without EdU/TuJ1 signal. Data are mean ± s.e.m. n = 6 (E9.5), n = 4 (E14.5). Kruskal-Wallis test with Dunn’s multiple comparisons test. Scale bars: 10 μm (e, f).

To identify novel signalling pathways regulating neuronal fate commitment of ENS progenitors advancing along the EPtG axis, we examined TrajectoryGeometry genes associated with the directionality of the neurogenic trajectory. Genes with positive scores are upregulated over the course of the trajectory, whereas those with negative scores are downregulated. As expected, genes with the highest positive values for neurogenesis had similar scores for the early and late neurogenic trajectories (for example *Elavl4, Pirt, Gap43*) (Fig. 2d). On the other hand, a subset of genes negatively associated with neurogenesis (such as *Top2a* and *Apoe*) exhibited divergent values between the early and late neurogenic trajectories, depending on their level of expression along the EPtG axis (Fig. 2d). We reasoned however that pathways critical for neuronal fate commitment across development, would be driven by genes exhibiting similar negative values for the early and late neurogenic branches, analogous to the profiles of *Sox10* and *Ednrb* (Fig. 2d), known regulators of ENS progenitor dynamics and differentiation^28^. Fulfilling these criteria was *Wwtr1* (Fig. 2d), which, together with *Yap1* (and its downstream target *Ccn1; Fig. 2d*), encode respectively the transcriptional co-activators TAZ and YAP of Hippo signalling, a pathway that regulates early neural crest delamination and lineage development^29, 30^. Immunostaining confirmed expression of YAP and TAZ in Sox10^+^ cells in the gut of mouse embryos (Fig. 2e). To test our prediction that Hippo signalling is implicated in enteric neurogenesis at different stages of ENS development, we used the Sox10CreERT2 driver^10^ to delete *Yap1* and *Wwtr1*^31^ from Sox10^+^ neural crest-derived cells in the gut of E9.5 and E14.5 mouse embryos. Single or combined conditional ablation of these genes reduced proliferation and expression of early neuronal markers (such as TuJ1) in the targeted cells within 48 hours (Fig. 2f), suggesting that Hippo signalling is required for both the expansion of undifferentiated ENS progenitors and their neuronal fate commitment at different stages along the EPtG axis. Together, our experiments argue that shared transcriptional mechanisms control neuronal commitment and differentiation of ENS progenitors throughout prenatal development, the period during which the vast majority of enteric neurons are generated^10^. Therefore, the neuronal diversity observed in the adult ENS is unlikely to be strictly linked to the timing of cell cycle exit but rather results from signalling events at postmitotic and relatively late stages of neuronal differentiation.

### Gene expression dynamics along the gliogenic axis

To uncover the regulatory basis for the transition of ENS progenitors from a neurogenic to gliogenic character, we first characterised ANTLER GMs exhibiting stage-specific expression along the EPtG axis. Consistent with the strong neurogenic output of eEPs (Fig. 1d), the top gene ontology term categories for GM16, 17 and 18, which were highly and selectively expressed by this cluster and their eENP progenies, were associated with the regulation of neuronal development (Fig. 3a, b; Extended Data Fig. 3). Included in these GMs were genes linked to WNT signalling (*Dvl3*, *Gsk3b*, *Fzd3*, *Peg12, Znrf3 and Bcat1*), a pathway that promotes CNS and ENS neurogenesis^32–35^ (Fig. 3a; for validations of expression see Extended Data Fig. 5a-c; Suppl. Table 1). Also included in these GMs were *Phox2b*, which is essential for autonomic neurogenesis^36^, *Dpysl2*, a regulator of axonal growth and guidance^37^, and *Hmga2* and its target *Igf2bp2*, which control stage-specific neural stem cell activity and enteric neuron development^38, 39^(Fig. 3a; for validations of expression see Extended Data Fig. 5d-f). Despite the widespread expression of neurogenic GMs in eEPs, our previous clonal analysis demonstrated that the majority (∼60 %) of Sox10^+^ cells in the gut of E12.5 mouse embryos give rise exclusively to glial cells^11^. In that respect, it was interesting that GM16 also included *Csde1*, an RNA binding protein that inhibits neurogenesis in a human embryonic stem cell differentiation model and was enriched in several of its known target transcripts^40–42^ (Suppl. Tables 3-6; for validations of expression see Extended Data Fig. 5g). We suggest therefore that the effective neuronal differentiation output of eEPs is controlled by the integrated activity of extracellular signals and transcriptional programs that drive neurogenesis, and posttranscriptional checkpoints that constrain neuronal differentiation to allow for the necessary expansion of the initially small ENS progenitor pool.

**Figure 3.**
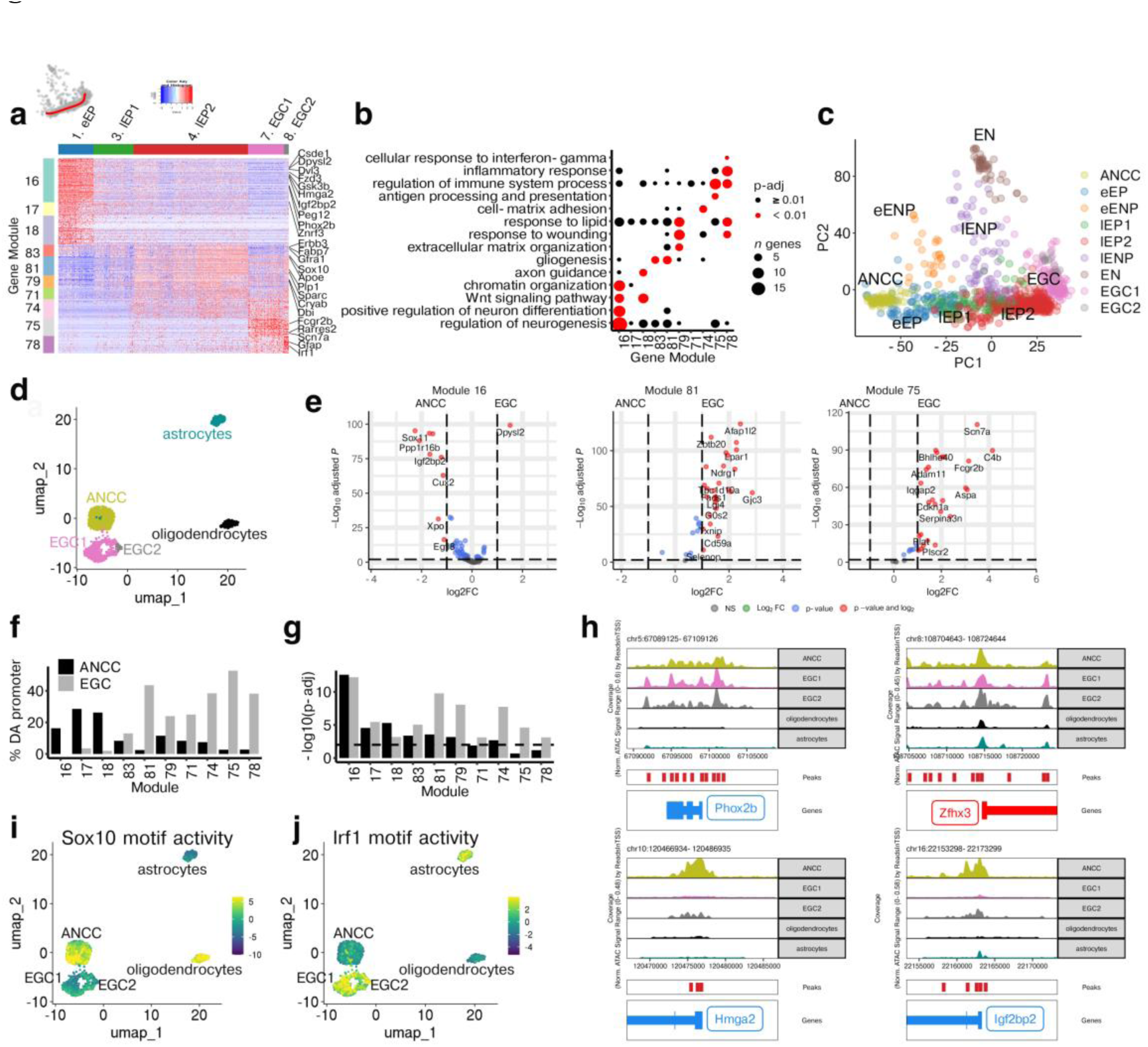
Transcriptional and epigenetic changes along the gliogenic trajectory. **a,** Heatmap (scaled normalised expression) for selected gene modules (left) associated with eEPs, lEP1/2, and EGC1/2 (top). Representative genes are indicated on the right. Inset indicates the gliogenic trajectory on the PCA. **b,** Dot plot showing statistical significance (colour of dot) and size of overlap (size of dot) between selected gene modules and indicated GO terms. **c,** PCA plot showing scRNA-seq datasets including tdT^+^ ENS cells (Fig. 1b,e) and ANCCs^3^. **d,** UMAP representation of scATAC-seq for EGCs, ANCCs, cortical astrocytes and cortical oligodendrocytes. **e,** Volcano plots showing mean log_2_-transformed fold change (FC; *x* axis) and significance (-Log_10_(adjusted P value)) of differentially accessible (DA) genes from the indicated GMs between ANCCs and EGCs. **f,** Bar plot indicating the percentage of genes per GM that have at least one DA peak in their promoter region between ANCCs and EGCs. **g,** Bar plot showing the statistical significance (hypergeometric test, -log_10_(p-adj)) of enrichment of genes with at least one peak in their promoter region for the indicated GMs. Dashed line indicates p-adj = 0.01. **h,** Track plots of ATAC signals for genes maintaining accessibility in EGCs (*Phox2b*, *Zfhx3*) and genes with reduced accessibility in EGCs (*Hmga2*, *Igfbp2*). **i,j,** UMAPs as in panel **d** indicating SOX10 (**i**) and IRF1 (**j**) motif activity, calculated using Chromvar.

To determine whether the neurogenic bias of eEPs is the legacy of pre-enteric neural crest progenitors^43, 44^ or a property acquired within the microenvironment of the embryonic gut, we performed a combined analysis of the transcriptome of ANCCs^3^ and our scRNA-seq datasets. ANCCs expressed the eEP/eENP-specific gene modules GM16-18 and principal component analysis positioned these cells at the start of the EPtG trajectory (Fig. 3c; Extended Data Fig. 6a), suggesting that ANCCs and eEPs share transcriptional profiles. Interestingly, GM16-18 were also expressed by diverse neural crest cell populations^3^ (Extended Data Fig. 6b), indicating that the transcriptional programmes underpinning the neurogenic potential of ANCCs and eEPs are expressed widely by neural crest cell lineages. A notable exception was *Phox2b*, which was specifically and uniformly induced in ANCCs and maintained at high levels in eEPs prior to its downregulation in lEPs and EGCs (Extended Data Fig. 6b, c; for validation of expression see Extended Data Fig. 5d, e). Interestingly, despite the strong neurogenic bias of ANCCs, no committed neuronal progenitors appeared to emerge from this cell population (Fig. 3c). A small number of GM16-18 genes, such as *Cux2*, *Egfl8*, *Runx1t1* and *Dlx5*, which have been implicated in mammalian neurogenesis, were specifically upregulated in eEPs (Extended Data Fig. 6d), suggesting that they play a role in converting the neurogenic bias of ANCCs into effective neurogenic output. Thus, our analysis indicates that ANCCs represent the incipient cell population of the EPtG trajectory and suggests that the neurogenic bias of eEPs is an intrinsic property of the autonomic lineage, which is licensed to undergo neurogenesis by the sequential induction of a small number of transcriptional regulators.

Relative to ANCCs/eEPs, lEPs downregulated the neurogenic modules GM16-18 and upregulated GMs (such as GMs 83, 81 and 79) which were related to glial development and the communication of cells with their tissue environment (Fig. 3a, b; Extended Data Fig. 3). Genes upregulated at this stage included known markers and regulators of gliogenesis (*Sox10, Apoe, Erbb3, Fabp7, Plp1*)^45–49^, extracellular matrix organization (*Sparc*)^50^ and response to wounding (*Cryab*)^51^ and also genes implicated in long-chain fatty acid metabolism in CNS glia (*Dbi*)^52^ (Fig. 3a, b; for validations of expression see Extended Data Fig. 5 h-k). Finally, in agreement with the emerging roles of enteric glia in gut immunity and host defence^7, 8^, GMs upregulated specifically by EGCs (GMs 71, 75, 78) included genes associated with the regulation of immune function and inflammatory responses (e.g *Fcgrt, Scn7a, Ifi27l2a, Rarres2, Serping1, Cxcl1, Irf1*)^53–59^ (Fig. 3a, b; for validations of expression see Extended Data Fig. 5l). Together, our analyses demonstrate that the transcriptional landscape of the EPtG trajectory is highly dynamic and changes according to the emerging anatomical and functional requirements of the gut: as development proceeds, the initially strong neurogenic character of Sox10^+^ cells, which ensures the efficient generation of enteric neurons, is replaced by cellular profiles befitting the emerging roles of this lineage in supporting the nascent intestinal neural circuits and maintaining immune and tissue homeostasis.

### Distinct chromatin accessibility patterns underpin the differential gene expression profile of ANCCs and EGCs

To examine whether chromatin remodelling is implicated in cell state transitions along the EPtG trajectory, we performed single-cell assay for transposase accessible chromatin using sequencing (scATAC-seq)^60^ on ANCCs and EGCs. UMAP visualisation showed that the two cell populations were clearly separated from each other (and from astrocytes and oligodendrocytes), indicating distinct chromatin accessibility profiles of these cell populations (Fig. 3d). Genes with promoter regions more accessible in ANCCs were associated with gene ontology terms related to neural development, tissue morphogenesis and cell fate commitment. In contrast, genes with promoter regions more accessible in EGCs were associated with terms related to cytokine responses and cell migration (Extended Data Fig. 7a). Therefore, in general, the transcriptional profiles of ANNCs and EGCs are concordant with their chromatin accessibility.

Since gene expression dynamics define cell state transitions along the EPtG axis, we next analysed the accessibility of genes from the time dependent GMs (Fig. 3a). As expected, genes for GMs characteristic of eEPs (GM16-18) or lEPs and EGCs (GMs 83, 81, 79, 71, 74, 75, 78) were more accessible in ANCCs or EGCs, respectively (Fig. 3e, Extended Data Fig. 7b). However, when considering the accessibility of individual peaks, it became apparent that for GM16 genes only very few of the differentially accessible (DA) peaks mapped to promoter regions (16.3 % of genes had promoter peaks more accessible in ANCCs, 0 % in EGCs), with the majority corresponding to intronic or distal elements (Fig. 3f; Extended Data Fig. 7c). Specifically, in EGCs 81.5 % of GM16 genes (including *Phox2b* and *Zfhx3*) contained at least one peak in their promoter region (Extended Data Fig. 7d), significantly more than expected by chance (Fig. 3g; hypergeometric test, p-adj = 6.2E-13, see materials and methods for details), indicating that despite the downregulation of this module along the EPtG axis, the majority of genes maintain an open chromatin configuration in their promoter. The remaining (18.5%) GM16 genes (which included the chromatin remodelling factor *Hmga2* and its target *Igf2bp2*) lacked promoter peaks in EGCs (Fig. 3f, h), suggesting that promoter accessibility of a subset of these genes regulates the neurogenic output of Sox10^+^ cells along the EPtG trajectory. Interestingly, the promoters of the handful of GM16-18 genes specifically upregulated in eEPs (*Cux2*, *Egfl8, Dlx5* and *Runx1tl*) were also accessible in ANCCs (Extended Data Fig. 7e), providing further evidence that the founder ENS cell population is primed for neuronal differentiation. Similarly, ANCCs had more GM83, 81, 79 genes (82.6 %, 76.9 %, 80 %) with peaks in their promoters than expected by chance (GM83 p-adj = 4.17E-4, GM81 p-adj = 2.76E-4, GM 79 p-adj = 6.67E-4) (Fig. 3g), suggesting that to some extent the chromatin organisation of these cells is prepared for the upcoming gliogenic gene expression profile. In contrast, for GM75 and 78, only EGCs had significantly more genes (77.8 %, 70.6 %) containing at least one peak in the promoter region than expected by chance (GM75 p-adj = 2.45E-5, GM75 p-adj = 7.51E-4) (Fig. 3g), arguing that cues from the gut environment are necessary for the chromatin remodelling associated with immune gene expression and the mature EGC phenotype.

To explore further the regulatory roles of the ANCC and EGC chromatin landscapes on gene expression, we searched for transcription factor motifs enriched in differentially accessible chromatin regions. Among the top hits in ANCCs were motifs for known transcriptional regulators of neural crest cell and ENS development (TFAP2a, TFAP2c and SOX10)^9, 61^ (Extended Data Fig. 7f; Supplementary Table 7). Interestingly, despite EGCs maintaining high levels of *Sox10* expression (Extended Data Fig. 7g), SOX10 motif activity was reduced in these cells (Fig. 3i), suggesting that stage-specific utilisation of transcription factor binding sites contributes to the functional specialisations of cell states along the EPtG trajectory. In contrast to ANCCs, EGC chromatin peaks were enriched in motifs for the AP1 family members FOS and JUN (Extended Data Fig. 7f; Supplementary Table 8), mediators of diverse cellular responses^62^. Consistent with the regulatory roles of the IFNγ-EGC signalling axis^8^, accessible chromatin in enteric glia was also enriched in motifs for interferon regulatory factors (IRFs)^63^ (Fig. 3j, Extended Data Fig. 7f). Taken together, our experiments indicate that chromatin remodelling of Sox10^+^ cells along the EPtG trajectory regulates the transition of ANCCs from effective neurogenic progenitors to enteric glia, which in addition to their canonical neuroprotective roles, regulate intestinal immune responses and tissue homeostasis. Nevertheless, our data reveal that mature EGCs maintain an epigenetic memory of their neurogenic past.

### Neurogenic potential of EGCs

Our model of ENS lineage configuration (Fig. 1e, f) and chromatin accessibility analysis is consistent with previous studies^10, 17, 18, 64^ and suggests that in certain tissue contexts, EGCs can return to upstream positions of the EPtG trajectory and re-activate neurogenic programs that are employed by early ENS progenitors *in vivo*. To examine this, we developed a robust cell culture model of EGC activation and differentiation (see materials and methods for details). Within 4 days after plating (days in vitro-DIV) lineally marked (tdT+) EGCs acquire the morphology of early neural crest cells and re-enter the cell cycle (Fig. 4a,b). Bulk and scRNA-seq datasets show that DIV4 cultures downregulate glial markers (*S100b*) and genes expressed by mature enteric glia (GM81 and GM75) and upregulate cell cycle markers (e.g. *Mki67*) and genes expressed by ANCCs and eEPs (GM16) (Fig. 4g,h; Extended Data Fig. 8a-d, f-h). By DIV11, cultures upregulate the neurogenic genes *Ret* and *Ascl1* expressed by early ENS progenitors (Extended Data Fig. 8d, e) and by DIV20 they form interconnected clusters reminiscent of in vivo ganglionic networks (called “ganglioids”) (Fig. 4c). Immunostaining confirms that neurons in ganglioids differentiate from tdT^+^ glia and express markers of synaptogenesis (SYN1) and markers of common neuronal subtypes in the ENS, such as NOS, VIP, CALB, NPY (Fig. 4d,e). Furthermore, our scRNA-seq analysis shows that the neuronal clusters of ganglioid cultures express key neurogenic markers and GMs characteristic of *in vivo* neurogenesis (Fig. 4i; Extended Data Fig. 8i). Using CRISPR-Cas9-mediated gene editing and pharmacological inhibition, we also demonstrated that the key regulators of enteric neurogenesis *Ret* and *Ascl1* were required for neuronal differentiation of ganglioids (Fig. 4j, Extended Data Fig. 8j-n). In addition, and similar to the role of *Foxd3* in maintaining multipotency and neurogenic potential of neural crest cells *in vivo*^65, 66^, we found that ganglioid cultures established from *Sox10CreERT2;Foxd3^fl/fl^* mice (see Materials and Methods) had a reduced number of neurons (Fig. 4k, Extended Data Fig. 8o, p). Finally, patch clamp analysis showed that retinoic acid, which promotes enteric neuron development^67^, enhanced the differentiation and functional maturation of EGC-derived enteric neurons in ganglioid cultures (Fig. 4f, l, m). Together, our experiments demonstrate that mammalian EGCs are capable of reactivating neurogenic programmes operating in early ENS progenitors and support the view that the EPtG trajectory of the ENS constitutes a transcriptomic continuum occupied by phenotypically adaptable cell states whose functions are linked to the emerging developmental and physiological requirements of the gut.

**Figure 4.**
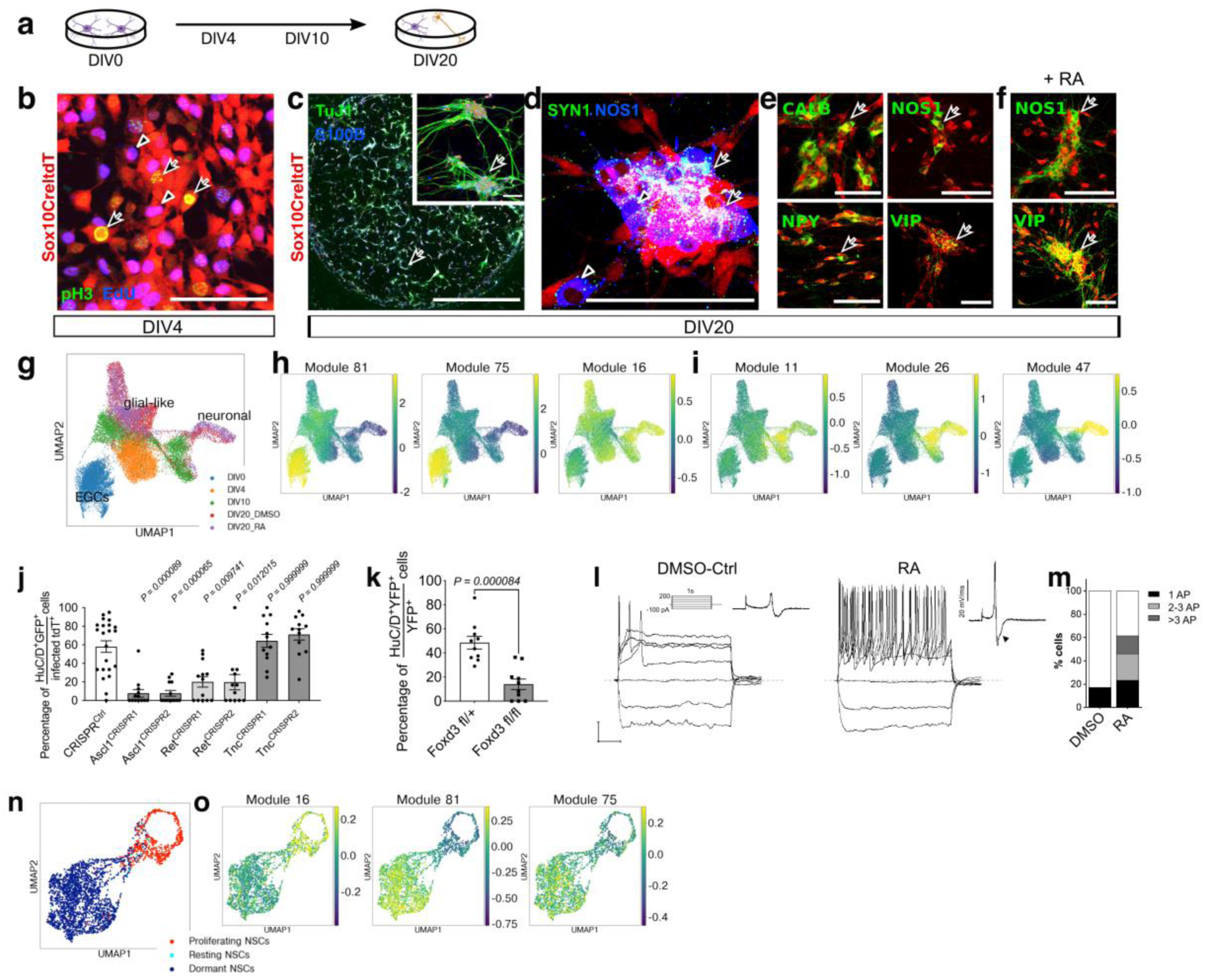
Ganglioid cultures recapitulate key molecular and functional characteristics of enteric neurogenesis. **a,** Schematic indicating the key time points ganglioid cultures were analysed. **b,** Ganglioid cultures at DIV4 indicating that cells derived from tdT^+^ EGCs (red) incorporate EdU (blue, arrowhead) and are labeled by pH3 (green/yellow, arrow). **c-e,** Immunostaining of ganglioid cultures with TuJ1 (green) and S100B (blue) (arrows indicate ganglioids) (**c**), SYN1 (green, arrows) and NOS1 (blue, arrowheads) (**d**), CALB, NOS1, NPY and VIP (green, arrows) at DIV20 (**e**). Scale bars: 5 cm (**c**), 500 μm (**c**, insert), 100 μm (**d**, **e**). **f,** Immunostaining of retinoic acid-supplemented ganglioid cultures for NOS1 (green, arrows, top) and VIP (green, arrows, bottom) at DIV20. Scale bars: 100 μm. **g,** UMAP representation of sequenced cells from ganglioid cultures colour-coded according to DIV. **h, i,** Expression (score) of developmental time (**h**) and enteric neurogenesis (**i**) -associated GMs on the UMAP shown in **g**. **j,** Quantification of neurons following CRISPR editing of ganglioid cultures at DIV20. Data are mean ± s.e.m. (n = 22 (CRISPR^CTRL^), n = 13 (Ascl1^CRISPR1+2^, Ret^CRISPR1^), n = 12 (Ret^CRISPR2^, Tnc^CRISPR1+2,^) fields of view per group). Kruskal-Wallis test with Dunn’s multiple comparisons test. **k,** Quantification of neurons in cultures from *Sox10CreER^T^*^2^*;Foxd3^fl/+^* and *Foxd3^fl/fl^* mice at DIV20. Data are mean ± s.e.m. (n= 10, field of view per group, pooled from two independent experiments). Unpaired student’s t-test. **l,** Superimposed membrane voltage responses of a representative action potential-generating DMSO (left) or retinoic acid-treated (right) neuron following 1s current injections from −100 to +200 pA in 50 pA increments (scale bars: horizontal 200 ms, vertical 20 mV). Corresponding time derivative waveforms of action potentials are shown as inserts. Arrowhead points to the presence of a hump, which is a defining feature of AH-type enteric neurons (control 0/5 = 0%; RA 14/15 = 93.3%). **m,** Bar plot showing proportion of cells firing action potentials in each treatment group: control (untreated and DMSO-ctrl cells) n= 29; RA treated: n= 26. **n,** UMAP representation of sequenced dormant (quiescent), resting and proliferating (active) hippocampal NSCs^71^. **o,** Expression (score) of the indicated GMs on the UMAP shown in **n**.

Next we considered the possibility that cell state transitions along the EPtG axis (during development and in ganglioid cultures) are analogous to adult neural stem cells switching between activated and quiescent states^68^. In support of this idea, GM16 (which is highly expressed by eEPs) and GMs 81, 71 and 74 (prominent in lEPs and EGCs) were enriched for genes upregulated in activated and quiescent neural stem cells, respectively^69, 70^ (Suppl. Tables 9, 10). Furthermore, GM16 showed highest expression in a scRNA-seq dataset of activated hippocampal neural stem cells^71^, whereas the corresponding quiescent neural stem cell population expressed modules characteristic of lEPs and EGCs (e.g. GMs 81 and 75) (Fig. 4n,o). Therefore, we suggest that common transcriptional mechanisms are implicated in the regulation of cell state transitions of neural progenitors in both the CNS and PNS.

## Discussion

In this study we have examined how the differentiation trajectories of enteric neurons and glia emerge during mammalian development. Our analysis supports a model according to which neurogenic differentiation paths branch from a linear gliogenic trajectory that traces the founder SOX10^+^ cell population as it loses progressively its pro-neurogenic and proliferative biases and acquires features of quiescent enteric glia. The direct anchoring of postmitotic neurogenic trajectories on the gliogenic axis and the positive association between neurogenic output and proliferation of progenitor cells, ensures that the overall size and balance of the neuronal and glial cell populations in the gut are regulated effectively by the cell cycle dynamics of SOX10^+^ progenitors. Our model also provides an explanation for the long-standing observation that the majority of mutations implicated in Hirschsprung’s disease, a relatively common neurodevelopmental abnormality of the ENS, have been identified in genes critical for the proliferation of ENS progenitors and lead to the concomitant loss of enteric neurons and glia^72^.

In contrast to the egress of neurogenic trajectories, which is associated with a major shift in the direction of change in gene expression, cells along the gliogenic axis maintain a core transcriptional profile established in early ENS progenitors, along with time dependent changes that allow them to adjust to the shifting anatomical and functional requirements of the host gut. Thus, as their initial neurogenic tasks are completed, ENS progenitors activate gene modules that allow them to support the nascent neural circuits and participate in tissue homeostasis and host defence. However, mature EGCs are capable of returning to developmentally upstream positions of the gliogenic axis and re-activating neurogenic programs. This occurs in certain contexts, such as ganglioid cultures of mammalian EGCs or following injury of the mouse gut, whereas EGCs in the adult zebrafish are capable of undergoing constitutive neurogenesis at steady state^10, 17^. Our findings indicate that the potential to return to neurogenic states may be facilitated by mature EGCs maintaining the expression of, and redeploying into new roles, central regulators of enteric neurogenesis (*Sox10*), or preserving an open chromatin configuration of their loci (*Phox2b*). We suggest that the gliogenic trajectory of the mammalian ENS represents a continuum occupied by dynamic transcriptional states that correspond to hierarchies of roles assumed by SOX10^+^ cells during intestinal organogenesis and within the tissue and luminal environment of the postnatal gut. Finally, our work has uncovered previously unappreciated analogies between cell state transitions along the gliogenic axis of the ENS during development and the cellular states defined by activated and quiescent neural stem cells in the adult brain. Characterisation of the cell-intrinsic mechanisms and environmental signals that allow EGCs to re-activate their neurogenic potential could have far reaching implications for the development of therapies aiming at restoring neural function compromised by injury or disease in the peripheral and central nervous system.

---

**Extended Data Fig. 1.**
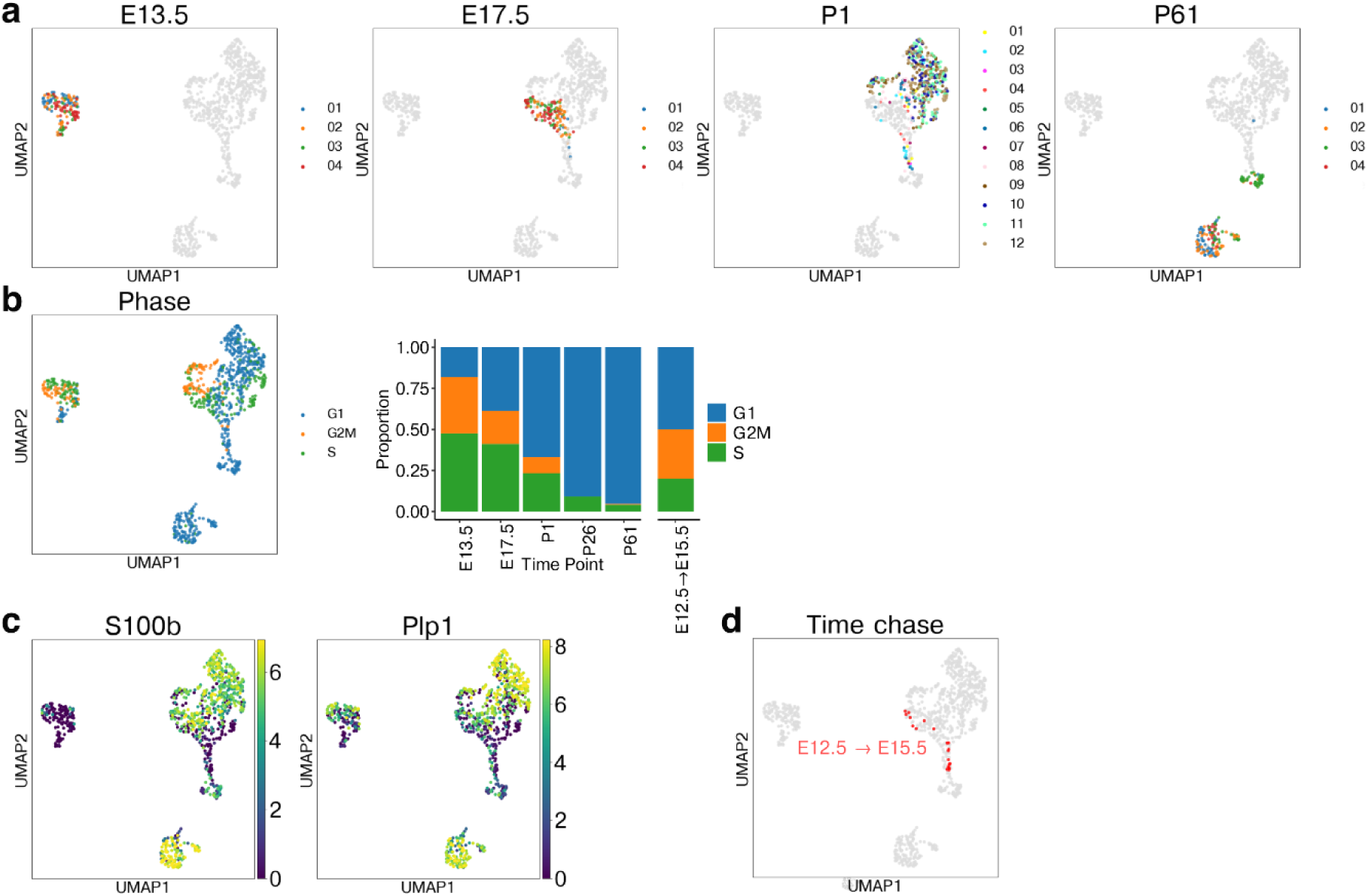
Properties of the transcriptional landscape of the developing ENS. **a,** UMAP representation of the scRNA-seq dataset with cells from an individual time point coloured by plate and cells from other time points coloured in grey. Note the lack of batch effects. **b,** UMAP representation of the entire scRNA-seq dataset coloured by inferred cell cycle phase and bar plot showing the proportions of inferred cell cycle phases at each time point. **c,** UMAPs coloured by expression of *S100b* and *Plp1*. **d)** UMAP showing lineage traced cells (red) labelled at E12.5 and harvested at E15.5.

**Extended Data Fig. 2.**
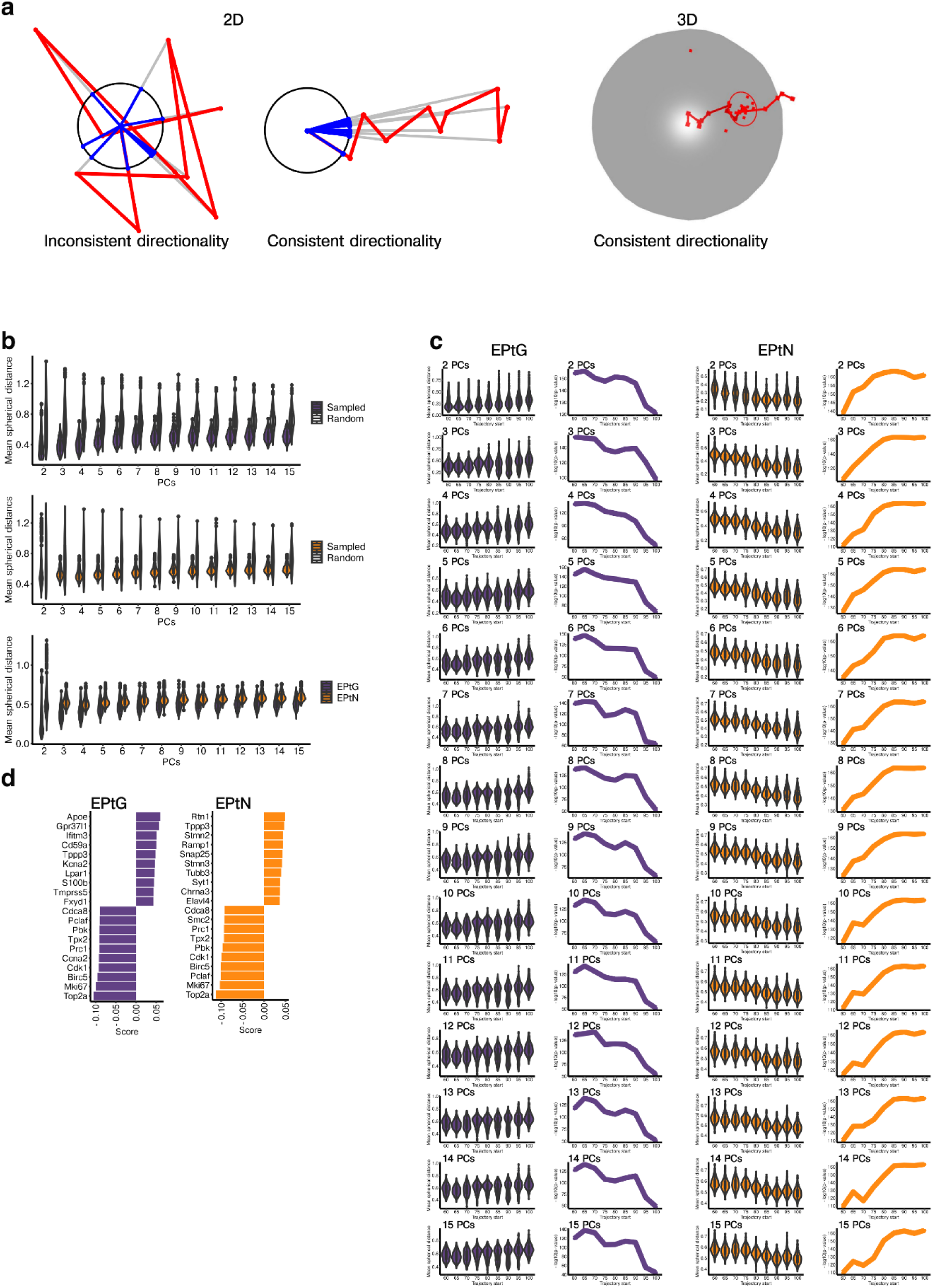
TrajectoryGeometry, a novel tool for the analysis of developmental trajectories. **a,** Illustration of the TrajectoryGeometry algorithm. A trajectory in 2 dimensions can be projected onto the 1-sphere (a circle). If this has inconsistent directionality the projections will disperse around the circumference, whereas if the trajectory has a consistent directionality these will be restricted to a small region. This approach can be extended to spheres in an arbitrary number of dimensions e.g. a trajectory in 3 dimensions can be projected onto the 2-sphere. Here the closeness of projections on the surface of the sphere can be described by a circle that shows the mean spherical distance from the centre of the projections. **b,** Violin plots showing the mean spherical distance (radii of the circles as shown in **a**) for paths sampled from glial and neuronal trajectories in comparison to random trajectories, and directly comparing glial and neuronal trajectories, when considering different numbers of PCs. **c,** Violin plots and line graphs showing –log10(p-value) for the significance of neuronal and glial trajectory directionality, relative to random trajectories, starting from successively later points in pseudotime, as the branch point is approached. **d,** Bar plots showing the top 10 TrajectoryGeometry genes positively associated (positive score) and top 10 TrajectoryGeometry genes negatively associated (negative score) with the directionality of the glial and neuronal trajectories.

**Extended Data Fig. 3.**
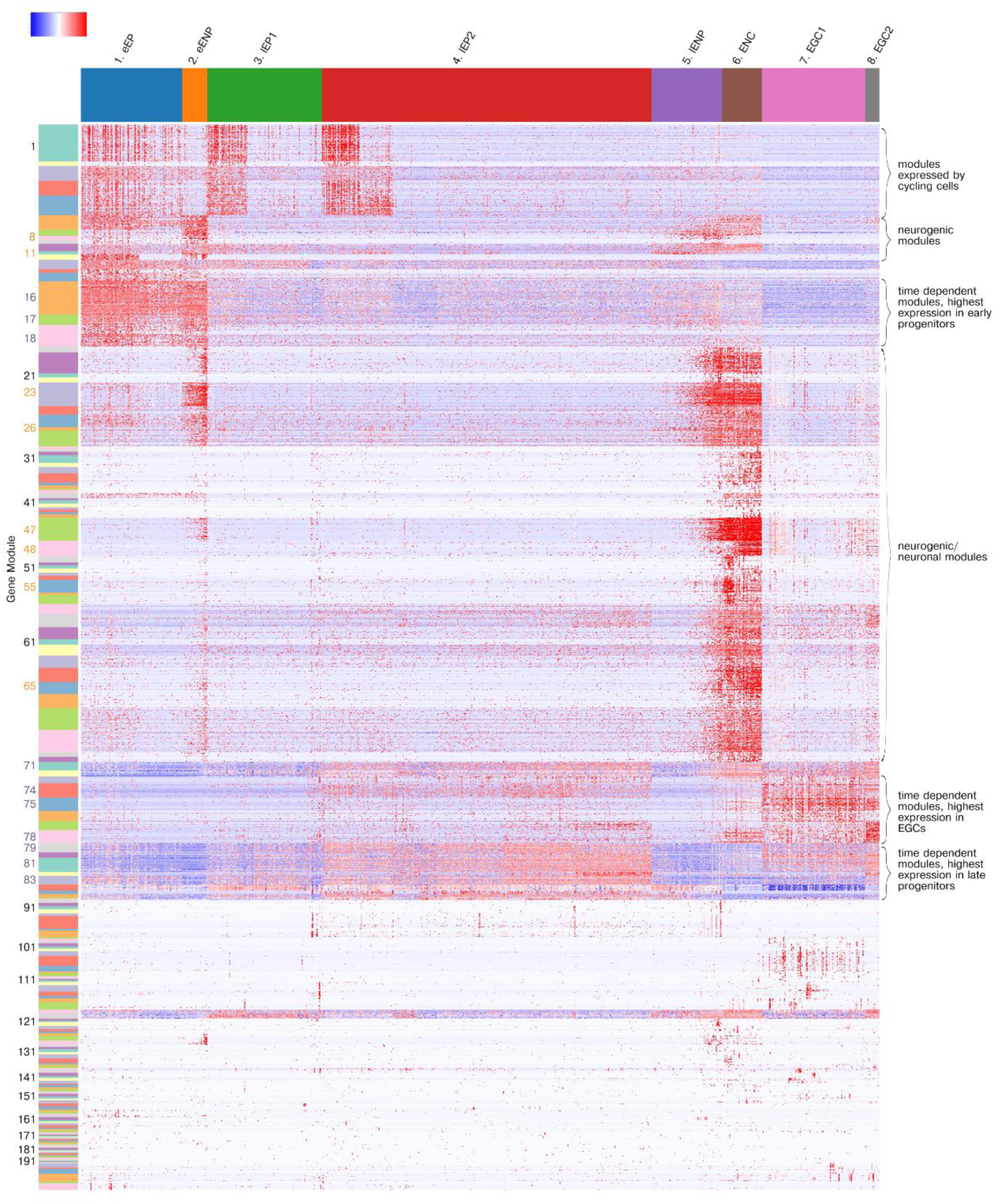
ANTLER gene modules. Heatmap (scaled normalised expression) of all gene modules (numbered on the left) identified using the ANTLER algorithm. Orange module numbers refer to neurogenic modules described in the main text; purple module numbers refer to modules with time-dependent expression described in the main text. Cell clusters are shown at the top.

**Extended Data Fig. 4.**
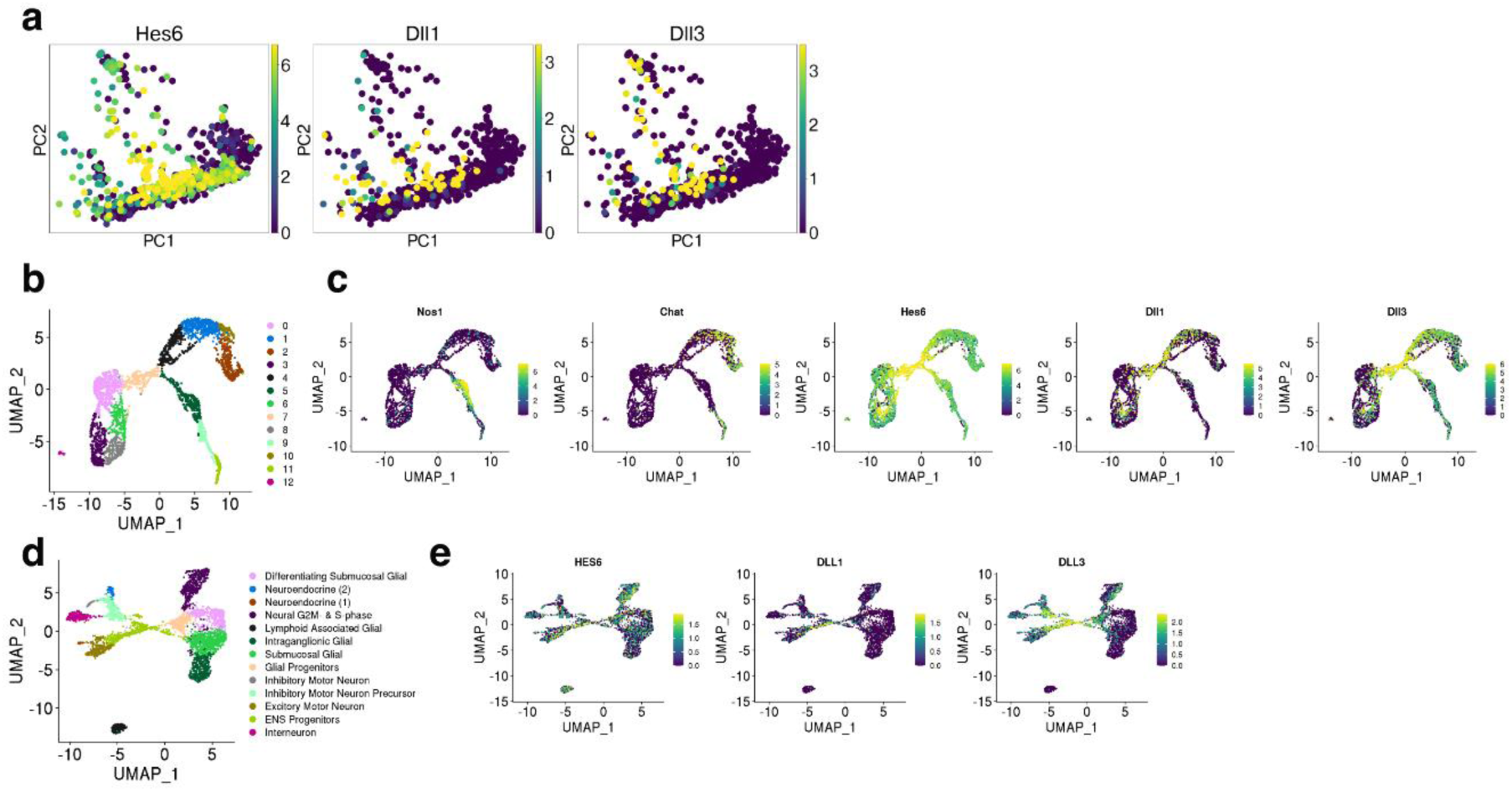
The expression of genes involved in Notch signalling. **a,** PCA plot (Fig. 1e) coloured by expression of *Hes6*, *Dll1* and *Dll3*. **b,** UMAP representation of E15.5 murine ENS cells^16^ coloured by Louvain clusters (resolution 1). **c,** UMAP as in **b** showing expression of *Nos1*, *Chat*, *Hes6*, *Dll1*, *Dll3*. **d,** UMAP representation of human ENS data (8-22 PCW)^14^. **e,** UMAP as in **d** showing expression of *HES6*, *DLL1*, *DLL3*.

**Extended Data Fig. 5.**
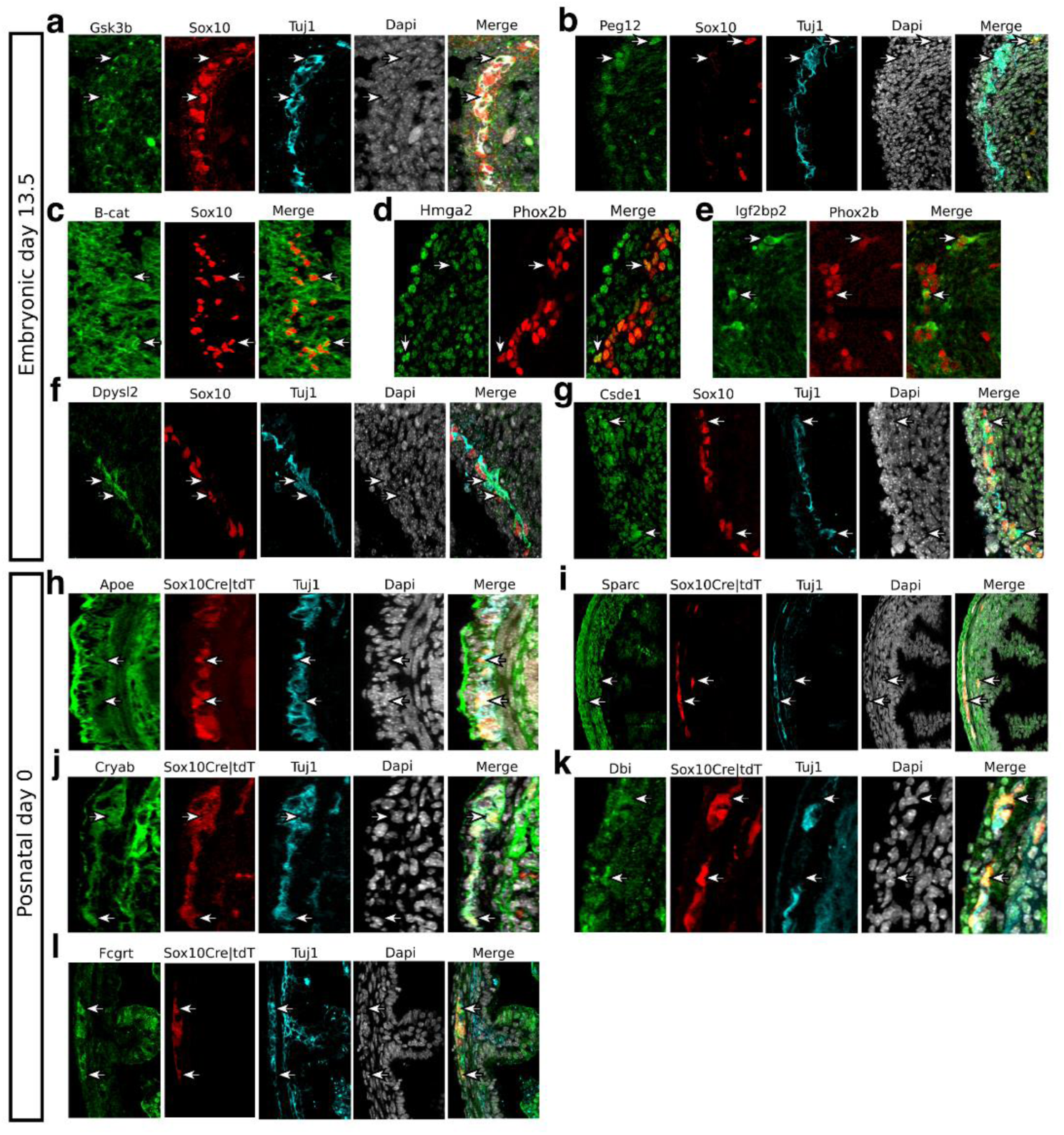
Validation of gene expression in the developing ENS. Immunostaining for the validation of expression of genes from ANTLER GMs in the ENS of mice at E13.5 (**a-g**) and P0 (**h-l**). Arrows point to cells of the ENS lineages.

**Extended Data Fig. 6.**
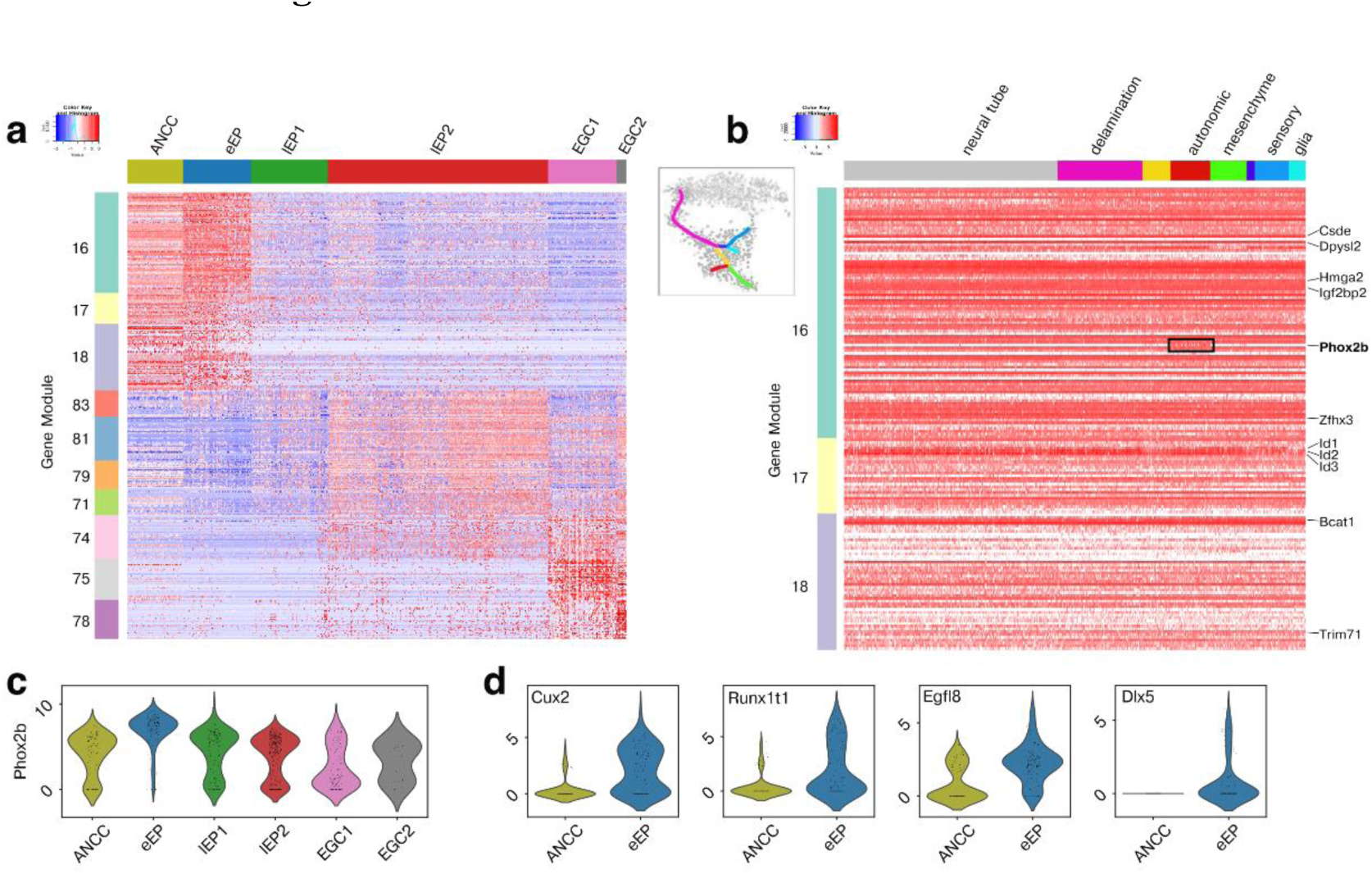
Neurogenic GM expression by early neural crest and ANCCs. **a,** Heatmap (scaled normalised expression) of the indicated GMs (left) in ANCCs^3^ and the tdT^+^ ENS cell clusters (top). **b,** Heatmap of gene expression (normalised, not scaled) for the neurogenic GM16-18 (left) in the indicated neural crest cell populations^3^. *Phox2b* expression is restricted to the autonomic lineage. **c,** Violin plots showing the normalised expression of *Phox2b* for ENS populations and ANCCs. **d,** Violin plots showing the normalised expression of *Cux2*, *Runx1tl*, *Egfl8* and *Dlx5* in ANCCs and eEPs.

**Extended Data Fig. 7.**
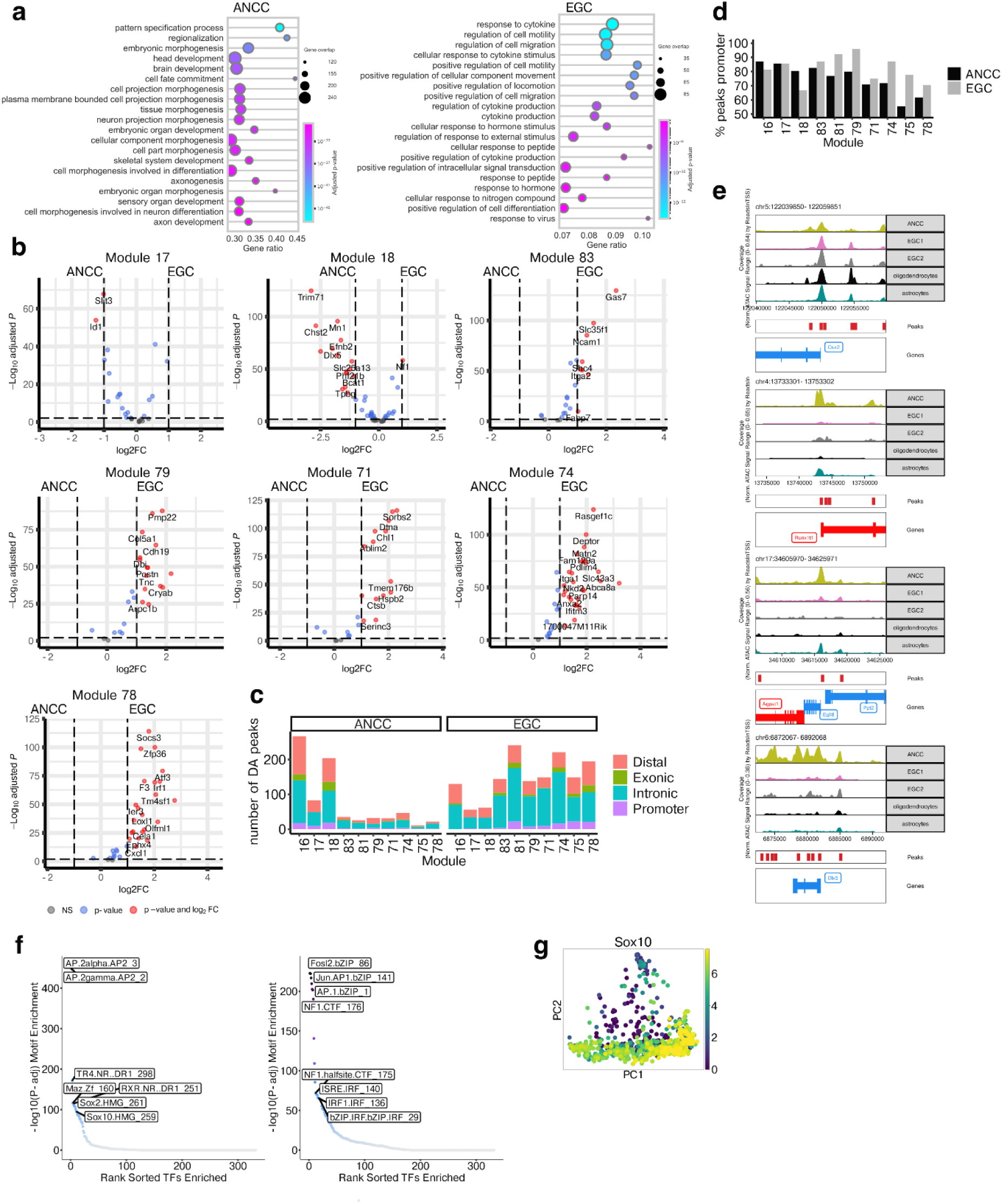
Chromatin accessibility in ANCCs and EGCs. **a,** Dot plot showing GO terms overrepresented among genes with differentially accessible peaks (log2FC > 1 & padj < 0.01) in promoter regions for ANCCs and EGCs. Dot size indicates the overlap for each term, and gene ratio indicates the fraction of genes in each term. **b,** Volcano plots showing differential accessibility between ANCCs and EGCs (based on gene score) for genes from the indicated GMs. **c,** Bar plot showing the number of DA peaks (log2FC > 1 & padj < 0.01) in ANCCs and EGCs. Results shown for peaks associated with genes from the indicated GMs. Colour indicates the gene region quantified (promoter, exonic, intronic, distal). **d,** Bar plot showing the percentage of genes in the indicated GMs that have at least one peak in their promoter region. **e,** Track plots showing gene accessibility for *Cux2*, *Runx1tl*, *Egfl8* and *Dlx5*. **f,** Plot showing enrichment of HOMER motifs in DA peaks (log2FC > 1 & padj < 0.01) in ANCCs and EGCs. **g,** PCA plot (Fig. 3c) showing expression of *Sox10*.

**Extended Data Fig. 8.**
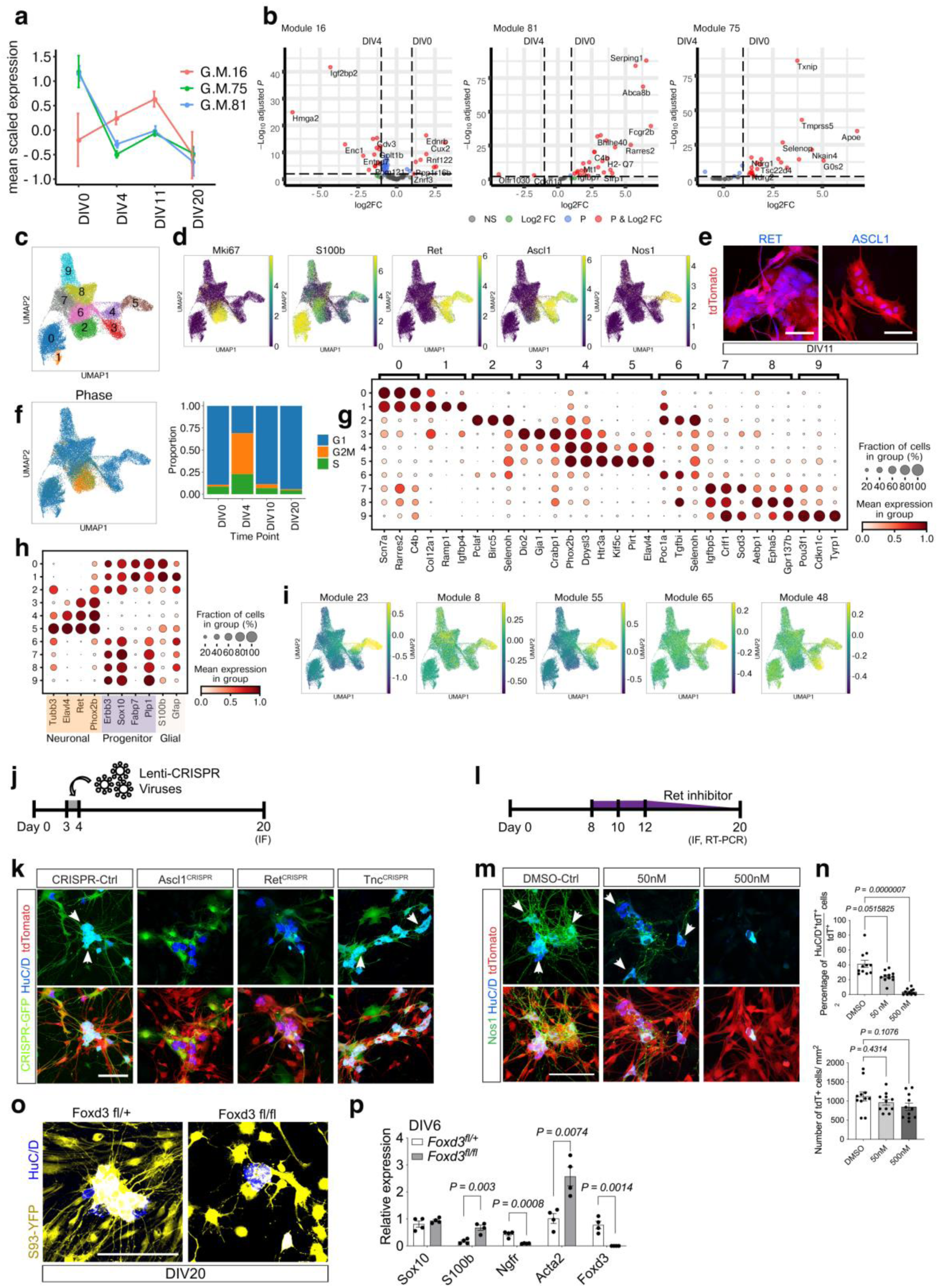
Enteric glial cell cultures recapitulate in vivo neurogenesis. **a,** Mean (± standard deviation) scaled expression of GM16, 75 and 81 in bulk RNA-seq datasets generated from the indicated DIV time points of ganglioid cultures. **b,** Volcano plots showing mean log_2_-transformed fold change (FC; *x* axis) and significance (-Log_10_(adjusted P value)) of differentially expressed genes from the indicated GMs at DIV0 and DIV4 of ganglioid cultures. **c,** UMAP representation of scRNA-seq data for ganglioid cultures coloured by Louvain clustering. **d,** Expression of the indicated genes on the UMAP in **c**. **e,** Ganglioid cultures (DIV11) immunostained for RET and ASCL1. **f,** UMAP shown in **c** coloured by inferred cell cycle phase and bar plot showing the proportions of inferred cell cycle phases at each time point in ganglioid cultures. **g,** Dot plot showing the expression of identified marker genes in the clusters shown in **c**. Dot size indicates the fraction of cells expressing an indicated marker while colour indicates mean expression of the gene marker. **h,** Dot plot showing the expression of neuronal, progenitor and glial markers in the clusters shown in **c**. **i,** Expression (score) of selected neurogenesis associated gene modules shown on the UMAP as in **c**. **j,** Experimental strategy for CRISPR editing of ganglioid cultures. **k,** CRISPR-virus infected ganglioid cultures at DIV20. tdTomato (red) indicates cells originating from tdT^+^ EGCs. GFP (green) and HuC/D (blue) identify CRISPR-infected and neurons (arrows), respectively. **l,** Experimental strategy for the pharmacological inhibition of RET signalling in ganglioid cultures. **m,** Ganglioid cultures (DIV20) immunostained for nNOS1 and HuC/D after inhibition of RET signalling. tdTomato (red) indicates cells originating from tdT^+^ EGCs. Arrows indicate neurons. **n,** Quantification of HuC/D+ cells upon RET signalling inhibition. Data are mean ± s.e.m. (n=11, field of view per group). Kruskal-Wallis test with Dunn’s multiple comparison test (top) and One-way ANOVA with Dunnett’s multiple comparison test (bottom). **o,** Representative images of ganglioid cultures derived from lineage-traced (YFP+, yellow) cells from *Sox10CreER^T^*^2^*;Foxd3^fl/+^*and *Sox10CreER^T^*^2^*;Foxd3^fl/fl^* mice. **p,** Relative gene expression of indicated genes in ganglioid cultures (DIV6) from *Sox10CreER^T^*^2^*;Foxd3^fl/+^* and *Sox10CreER^T^*^2^*;Foxd3^fl/fl^* mice. Data are mean ± s.e.m. (n = 4). Unpaired two-tailed t-test. Scale bars: 50 μm (e), 100 μm (k, m, o).

## Methods

### Data reporting

No statistical methods were used to predetermine sample size. Investigators were not blinded during experiments or outcome assessment.

### Animals

All animal procedures were carried out at the Francis Crick Institute in accordance with the regulatory standards of the UK Home Office (ASPA 1986) and the ARRIVE guidelines and approved by the local Animal Welfare and Ethics Review Body (AWERB). Mice were housed and bred under specific pathogen-free conditions (SPF) in individually ventilated cages under a 12 h light–dark cycle at ambient temperature (19 °C–21 °C) and humidity (45–55%). Standard food and water were provided ad libitum. Both male and female animals were used for the experiments. The *Sox10CreER^T^*^2^ transgenic mice refer to two sublines, SER26 (MGI: 5301107) and SER93 (MGI: 5910373) that have been reported previously^8, 10, 11^. Generation of the *Yap1^fl^* (MGI: 5603606), *Wwtr1^fl^* (MGI: 5603602) and *Foxd3^fl^* (MGI: 3790794) alleles and the *Wnt1Cre2* transgene have been described previously ^31, 66, 73^. The Cre-dependent reporters used are: *Rosa26-tdTomato* (MGI: 3809524)^74^, *Rosa26-nuclearGFP* (MGI: 5443817)^75^ and *Rosa26*-e*YFP*^76^. Sox10CreER|tdT and Sox10CreER|YFP indicate *Sox10CreER^T^*^2^*(SER93);Rosa26-tdTomato* and *Sox10CreER^T^*^2^*(SER93);Rosa26-YFP* mice, respectively. Mice of the desired developmental stage were generated by setting up timed pregnancies. The plug day was designated as E0.5 and date of birth as P0.

### Phenotypic characterisation of Yap1 and Wwtr1 deletion

For the *Yap1/Wwtr1* deletion experiments (Fig. 2f), the *Sox10CreER^T^*^2^ transgene was combined with the *Yap1^fl^* and *Wwtr1^fl^* alleles and the Cre-dependent *Rosa26*-e*YFP* reporter^76^. Gene deletion was induced by intraperitoneal injection of tamoxifen (Sigma, T-5648; stock solution 20 or 40 mg/ml made in corn oil/10% ethanol) to *Sox10CreER^T^*^2^*(SER93);Yap1^fl/fl^;Rosa26-eYFP* or *Sox10CreER^T^*^2^*;Wwtr1^fl/fl^;Rosa26-eYFP or Sox10CreER^T^*^2^*;Yap1^fl/fl^;Wwtr1^fl/fl^;Rosa26-eYFP* mice at a dose of 50 μg/g (for E9.5 embryos) or 100 μg/g (for E14.5 embryos) of dam body weight. For in vivo labelling of proliferating SOX10^+^ cells 5-Ethynyl-2’-deoxyuridine (EdU, 0.3 mg/ml, Thermo Fisher Scientific, E10187) was administered intraperitoneally to pregnant females two hours prior to harvesting of embryos. Embryonic guts were dissected 48 hours post tamoxifen injections (i.e. at E11.5 or E16.5), fixed overnight in ice-cold 4% paraformaldehyde and processed further for cryostat sectioning (10 µm). Briefly, samples were subjected to antigen retrieval using sodium citrate (10 mM) at pH 6.0, permeabilized with 0.3% Triton X-100 phosphate buffer (PBT), blocked with 10% donkey serum in PBT for an hour, incubated overnight with primary antibodies at 4 °C, rinsed at room temperature and further incubated with secondary antibodies for two hours at room temperature. After final washes, samples were processed for EdU labelling using the Click-iT EdU Alexa Fluor 647 EdU labelling kit following the manufacturer’s instructions (Thermo Fisher Scientific, #C10340). Slides were mounted for confocal imaging with Vectashield Mounting medium (Vector Labs, H-1000 /H-1200).

### Ganglioid cultures

Neurogenic EGC cultures were established from Sox10Cre|tdT mice 24 hours after tamoxifen injection (one dose at 100μg/g of body weight) and *Sox10CreERT2;Foxd3^fl/fl^;Rosa26-eYFP* mice 48 hours after tamoxifen injection (two doses of 100μg/g of body weight). Longitudinal muscle-myenteric preparations were washed thoroughly in DMEM/F12 media containing fetal bovine serum (FBS, 2%, Sigma-Aldrich, F7524), penicillin-streptomycin (100U/ml, Thermo Fisher Scientific) and 12.5mM HEPES (12.5mM, Thermo Fisher Scientific). Post-rinsing, tissues were minced using fine scissors and incubated in a shaker at 37°C for 30 min with Type 1 collagenase (0.5 mg/ml, Sigma Aldrich, SCR103) and DNaseI (0.5 mg/ml, Sigma-Aldrich, DN25) in the buffer described above. Next, trituration of the sample using an unpolished glass pasteur pipette was carried out. Triturated samples were then washed twice with DMEM/F12 media containing 10% FBS with penicillin-streptomycin and plated onto two 60 mm (Corning) dishes coated with fibronectin (20 μg/ml; Sigma-Aldrich, F1141). Seeded cells were cultured overnight in DMEM/F12 media containing 10% FBS and penicillin-streptomycin. Next day, the culture medium was changed to serum-free media that contained DMEM/F12 supplemented with N2 (1%, Thermo Fisher Scientific, 17502048), G5 (1%, Thermo Fisher Scientific, 17503012) and NGF-7S (50 ng/ml, Thermo Fisher Scientific, 13290-010). To allow robust proliferation of glial cells, half of the medium was replaced with fresh medium on day three. On day four, cells were trypsinized using TrypLE express (Thermo Fisher Scientific, 12604013) and filtered using a 40 μm cell strainer to obtain a single cell suspension. Cells were counted (Countess II; Thermo Fisher Scientific) and then seeded at ∼30,000 cells/cm^2^ onto fibronectin coated 60 mm dishes in proliferative medium. The passaged cells were collected using Accutase (Thermo Fisher Scientific, 00-4555-56) on day six and seeded onto Poly-D-Lysine/Laminin/Fibronectin coated 12 mm glass coverslips (Corning, #354087) at ∼42,000 cells/cm^2^ in 24-well plates. Upon passaging, cells were maintained in media that constitutes both the proliferative and neuronal differentiating medium which comprises of DMEM/F12, N2 supplement (1%), GDNF (10 ng/ml, Peprotech, 450-44), BDNF (10 ng/ml, R&D systems, 248-BD), NT3 (10 ng/ml, R&D systems, 267-N3), NGF-7S (10 ng/ml), cAMP (0.5 mM, Sigma-Aldrich, D0260), Ascorbic Acid (0.2 mM, Sigma-Aldrich, A4034) and penicillin-streptomycin at a ratio of 1:2. On day eight, the culture medium was completely replaced with neuronal differentiating media. Subsequently, every two days, half of the medium was exchanged with fresh media. To label proliferating cells, cultures were exposed to EdU (10 μM) for an hour before fixation. Cells were consistently maintained at 5% CO_2_.

For CRISPR editing, cultures were infected at DIV3 with lentiviruses carrying the Cas9 enzyme and sgRNA sequences to target indicated genes (Extended Data Fig. 8j). For RET signalling inhibition, indicated concentrations of GSK3179106, a selective RET kinase inhibitor^77^, or DMSO (control) were introduced on DIV8 and gradually reduced every two days from DIV12 onwards as shown in Extended Data Fig. 8l.

### CRISPR/Cas9-sgRNA design and lentivirus production

Seeded HEK293T cells were allowed to reach confluence in 10cm dishes with media containing DMEM with high glucose, GlutaMAX Supplement, pyruvate (Thermo Fisher Scientific, 10569010), FBS (10%) and penicillin-streptomycin. Next, cells were transfected with either a control pL-CRISPR.EFS.GFP (15µg, Addgene, 57818)^78^ or pL-CRISPR.EFS.GFP with cloned sgRNA sequences targeting Ascl1 (CRISPR1-F 5’CACCGCAACGAGCGCGAGCGCAACC-3’, CRISPR1-R 5’-AAACGGTTGCGCTCGCGCTCGTTGC-3’, CRISPR2-F 5’-CACCGGAGCATGTCCCCAACGGCG-3’, CRISPR2-R 5’-AAACCGCCGTTGGGGACATGCTCC-3’), Ret (CRISPR1-F 5’-CACCGAAGCGACGTCCGGCGCCGCA-3’, CRISPR1-R 5’-AAACTGCGGCGCCGGACGTCGCTTC-3’, CRISPR2-F 5’-CACCGTCTATGGCGTCTACCGTACA-3’, CRISPR2-R 5’-AAACTGTACGGTAGACGCCATAGAC-3’) and Tnc (CRISPR1-F 5’-CACCGTGTGACGATGGGTTCACAG-3’, CRISPR1-R 5’ AAACCTGTGAACCCATCGTCACAC-3’, CRISPR2-F 5’-CACCGCCCGCCGATTGTCACCACCG-3’, CRISPR2-R 5’-AAACCGGTGGTGACAATCGGCGGGC-3’). The constructs were provided with a packaging vector (Pax2, 12.5 µg) and the envelope protein VSV-G (3.3 µg, Addgene, #8454) in Opti-MEM medium to which the transfection reagent Polyethylenimine (1 mg/ml, Generon) was added at a ratio of 1:3 of the DNA. Guide RNA sequences were designed using Benchling [Biology Software] (2019; retrieved from https://benchling.com) and cloned based on a publicly available lentiviral CRISPR toolbox protocol^79^. Eight hours post-transfection, the supernatant was discarded and replaced with fresh medium. This was followed by collecting ∼10 ml of the filtered supernatant 48 hours later. The lentivirus was concentrated with the use of the PEG-it virus precipitation solution (2.5 ml, System Biosciences, LV810A-1) for 2-3 days at 4°C. Subsequently, precipitated viral particles were centrifuged at 2,300g for 30 min at 4°C, resuspended in proliferative media (250 μl) and stored at −80°C as 50 μl single-use aliquots for future use.

### Real time quantitative PCR of EGC cultures

Total RNA was isolated from FACS-enriched tdT+ cells of DIV20 using Trizol LS reagent (Thermo Fisher Scientific, 10296010) and the PureLink RNA Micro Kit (Thermo Fisher Scientific, 12183016) as per manufacturer’s instructions. Complementary DNA (cDNA) was generated by reverse transcription using the High-Capacity cDNA Reverse Transcription Kit (Thermo Fisher Scientific, 4368814). RT-qPCR was performed with the synthesized cDNA using Taqman fast universal 2X PCR Master Mix (Thermo Fisher Scientific, 4352042) and Taqman probes (Thermo Fisher Scientific) on a 7500 Fast Real-Time PCR system (Applied Biosystems). The Taqman probes used in this study are the following: *Actb* (Mm02619580_g1), *Acta2* (Mm00725412_s1), *Foxd3* (Mm02384867_s1), *Ngfr* (Mm00446296_m1), *Sox10* (Mm00569909_m1), *S100b* (Mm00485897_m1). C_t_ values obtained were normalized to β-actin and relative changes in expression were calculated using ΔΔC_t_ analysis.

### Bulk RNA sequencing of EGC cultures

Cultured cells were detached from the plates on DIV 4, 11 and 20 and enriched by FACS into collection tubes containing Trizol LS Reagent (Thermo Fisher Scientific, 10296010). Total RNA from 60,000 sorted cells/sample was isolated via the precipitation method in combination with spin-columns using PureLink RNA Micro Kit (Thermo Fisher Scientific, 12183016) according to the manufacturer’s instructions. cDNA was generated using the Ovation RNA-Seq System V2 (NuGen Technologies, 7102-A01) and quantified using the dsDNA Qubit HS Assay kit (Thermo Fisher Scientific, Q32854). This was followed by acoustic shearing of the cDNA to generate 200 bp fragments using Covaris E220 (Covaris). The Ovation Ultralow System V2 1-16 (NuGEN, 0344NB-A01) was used to produce libraries from fragmented double-stranded cDNA, followed by PCR amplification and library purification. The quality and quantity of the final libraries was assessed with TapeStation D1000 Assay (Agilent Technologies). The final libraries were pooled and sequenced on the HiSeq4000 (Illumina) to generate 75bp single-end reads.

### Single cell RNA sequencing

For the transcriptomic analysis described in Fig. 1, tdT^+^ cells were isolated at indicated stages from the small intestine of *Sox10CreER^T^*^2^*;Rosa26-tdTomato* (Sox10Cre|tdT) mice. For the isolation of E17.5, P1 and P26 cells, animals were injected with tamoxifen 20-24 hours earlier, while E15.5 cells were isolated from animals injected with tamoxifen at E12.5. The scRNA-seq datasets for E13.5 and P61 cells have been described previously^8, 11^. Expression of tdT was induced by intraperitoneal administration of tamoxifen to pregnant females at E12.5 and E16.5 and mice at P25 and P60 at 100 μg/g of body weight. For P1 cells, tamoxifen was injected intradermally at P0. Tissue was incubated with Collagenase I (1 mg/ml, Sigma, C0130) for 30-45 min and enzyme activity was terminated by rinsing cells with ice cold PBS. The cell suspension was centrifuged and filtered through 40 μm nylon cell strainers (Falcon, 352340). tdT^+^ cells were isolated by flow cytometry (BD Biosciences Aria Fusion sorter equipped with Diva V8 software) with a 100 μm nozzle and collected in 2% FBS in OptiMEM without phenol red (Thermo Fisher Scientific, 11058021). Dead cells were excluded by labelling with the cell viability dye Zombie Aqua (2 ng/ml, BioLegend, 423101) that was added to the single cell suspension before sorting. For the P1 stage, we also sorted the cells directly into 96-well plates to increase the yield. The protocols for the isolation of tdT^+^ cells from the small intestine of E13.5 embryos and P61 animals have been reported previously^8, 11^. RNA sequencing of isolated tdT^+^ cells was carried out as described previously^8, 11^. The Fluidigm C1 automated microfluidic system was used to capture individual cells into 5-10μm and 10-20μm chips. The SMARTer Ultralow RNA kit (Clontech, 634833) was used to reverse transcribe poly(A) RNA and amplify cDNA. ERCC spike-ins (Life Technologies, 4456740) were added at a dilution of 1:80000. cDNA concentration of each cell sample was quantified using the Promega Quantifluor dsDNA on Glomax system and quality of random samples was checked on high-sensitivity DNA chip on the Agilent Bioanalyser 2100. Amplified cDNA (0.125-0.375 ng) was used for generating sequencing libraries with the Illumina Nextera XT DNA kit (Illumina, FC-131-1096). All libraries were sequenced as paired-end 75 using the Illumina HiSeq4000 system.

For the transcriptomic analysis of adult EGCs (Fig. 4 and Extended Data Fig. 8), nGFP^+^ cells were isolated from tunica muscularis preparations from 8 week old *Sox10CreER^T^*^2^*(SER93);Rosa26-nuclearGFP* mice that had been injected with tamoxifen (2 doses of tamoxifen at 100 μg/g of body weight 10 days prior isolation). Tissue was incubated with 10 mg/ml collagenase IV (Worthington Biochemicals, CLS-4) in Hanks Balanced Salt Solution (HBSS, Thermo Fisher Scientific, 14170112) at 37 °C for 12 min. This was followed by a wash in ice-cold HBSS and subsequent incubation with papain (1U, Worthington Biochemicals, PAP2) for 5 min at 37 °C. The digested samples were then washed and resuspended in L-15 medium without phenol red (Thermo Fisher, 21083027) containing penicillin–streptomycin (1%, Thermo Fisher Scientific, 15140122), BSA (1 mg/ml, Sigma-Aldrich, A9418), HEPES pH 7.4 (10 mM, Thermo Fisher Scientific, 15630106), Biowhittaker water (10%, Lonza, BE17-724F) and DNase I (400U, Grade II; Roche, 10104159001), filtered through a 70 μm and 40 μm strainer and subjected to FACS using DAPI (1 μg/ml) for live/dead cell discrimination. For scRNA-seq of these cells we used the 10X genomics platform. Approximately 10,000 tdT^+^ cells post-FACS (from EGC cultures) and 10,000 nGFP^+^ cells post-FACS (freshly isolated EGCs) were loaded into a 10x Chromium Chip B (10x 3’v3) and partitioned into gel bead-in-emulsions (GEMs) using the 10x Chromium Controller. cDNA was generated following manufacturer’s protocol and quantified using the TapeStation D5000 ScreenTape (Agilent). Libraries were prepared according to the manufacturer’s guidelines and quality was monitored using both TapeStation (Agilent) and High Sensitivity dsDNA Qubit (Thermo Fisher Scientific) systems. Final libraries were pooled and sequenced using the HiSeq4000 (Illumina).

### Single-cell ATAC sequencing

For scATAC-seq of adult EGCs, nGFP^+^ cells were isolated from tunica muscularis preparations as described above. For ATAC sequencing of autonomic neural crest cells, tdT^+^ cells were isolated from *Wnt1Cre;Rosa26-tdTomato* embryos at E9.5 and E10.5. Cells were dissociated using mechanical and enzymatic disintegration as previously reported^80^. Briefly, heads were separated from trunks by slicing the embryos with a razor blade posteriorly to the otic vesicle, tissue was diced with a razor blade and triturated using a P-1000 pipette and incubated for roughly 15 min with Trypsin/EDTA at 37 °C. The enzymatic digestion was halted using ice-cold 1% FBS in HBSS. Cells were pelleted and filtered using a 40 µm cell strainer. Dissociated cell suspension was sorted (Tomato-positive from negative cells) using a FACSAria III flow cytometer. The sorted cells were stored in RMPI/DMSO buffer in −80 °C before nuclear isolation.

Nuclei were isolated and scATAC-seq libraries were prepared according to the Chromium Single Cell ATAC Reagent Kits User Guide (10x Genomics; 10xGenomics.com CG000496 Rev A). Briefly, after counting, nuclei concentrations were adjusted to the desired capture number 10,000 based on the number of available nuclei and the desired multiplet rate (described on 10X protocol). To minimize potential multiplets, we typically aimed to capture <6,000 nuclei per channel. Next, 5 μl of the resulting resuspended nuclei were combined with ATAC Buffer and ATAC Enzyme (10x Genomics PN-1000390) to form a transposition mix, which was then incubated for 60 min at 37 °C. A master mix composed of Barcoding Reagent, Reducing Agent B and Barcoding Enzyme (10X PN-1000390) was then added to the same tube as transposed nuclei. The resulting solution was loaded onto a Chromium Chip H(10x Genomics; PN-1000161). Vortexed Chromium Single Cell ATAC Gel Beads and Partitioning Oil were also loaded onto the same Chromium Chip H before attaching a 10x Gasket and placed into a Chromium Single Cell Controller instrument (10X genomics). Resulting single-cell GEMs were collected at the completion of the run (∼17 min) and linear amplification was performed on AB cycler: 72 °C for 5 min, 98 °C for 30 s, cycled 14×: 98 °C for 10 s, 59 °C for 30 s and 72 °C for 1 min. Emulsions were coalesced using the Recovery Agent (10x Genomics; 220016), then subjected to Dynabeads (2000048) and SPRIselect reagent (Beckman Coulter; B23318) bead clean-ups. Indexed sequencing libraries were constructed by combining the barcoded linear amplification product with a sample index PCR mix comprising SI-PCR Primer B, Amp Mix and Chromium i7 Sample Index Plate N, Set A (10x Genomics; 3000262). Amplification was performed on an AB cycler: 98 °C for 45 s, for 12 cycles: 98 °C for 20 s, 67 °C for 30 s, 72 °C for 20 s, with a final extension of 72 °C for 1 min. The sequencing libraries were subjected to a final bead clean-up SPRIselect reagent and quantified by Qubit and TapeStation HS1000. Sequencing libraries from adult EGCs were loaded on an Illumina Novaseq sequencer with 2 × 50 paired-end kits using the following read length: 51 bp read 1N, 8 bp i7 index, 24 bp i5 index (trimmed later on) and 51 bp read 2N using a loading molarity of 280 pM (Standard loading) or 180 pM (Xp loading).

### Sample preparation and Immunostaining experiments

Embryos (E9.5 - E13.5) and embryonic guts (>E16.5) were fixed overnight in ice-cold 4% paraformaldehyde (PFA) and processed further for cryostat sectioning (10 µM) or/and whole mount immunostainings. Briefly, samples were subjected to antigen retrieval using sodium citrate (10 mM) at pH 6.0, permeabilized with 0.3% Triton X-100 phosphate buffer (PBT), blocked with 10% donkey serum in PBT for an hour, incubated overnight with primary antibodies at 4 °C, rinsed at room temperature and further incubated with secondary antibodies for two hours at room temperature. After final washes, the samples were either processed for EdU labelling or mounted for confocal imaging with Vectashield Mounting medium (Vector Labs, H-1000 /H-1200). EdU labelling was done with Click-iT EdU Alexa Fluor 647 kit and the manufacturer’s instructions were followed (Thermo Fisher Scientific, #C10340).

*In vitro* cells cultured on cover slips and/or exposed to EdU (10µM) for one hour were fixed with 4% PFA at room temperature for 20 min. Fixed cells were rinsed with phosphate buffer, followed by antibody and EdU staining protocols as described above without the use of the antigen retrieval step. Stained coverslips were mounted onto glass slides using ProLong Diamond Antifade mountant (Thermo Fisher Scientific, P36961).

The following antibodies were used:GFP (Rat, Nacalai Tesque, #04404-84, 1:500), GFP (Chick, Abcam, #ab13970, 1:400), TuJ1 (Mouse, Biolegend, #801202, 1:1000), SOX10 (Rabbit, ProteinTech, #10422-1-AP, 1:200), S100 (Rabbit, DAKO, #z0311, 1:500), nNOS1 (Goat, Abcam, #ab1376, 1:300), HuC/D (Mouse, Thermo Fisher Scientific, #A-21271, 1:400), PHOX2B (Goat, R&D systems, #AF4940, 1:250), YAP1 (Mouse, Santa Cruz, #sc-376830, 1:200), WWTR1 (Rabbit, CST, #72804, 1:200), pH3 (Rabbit, Millipore, #06-570, 1:500), CALB1 (Rabbit, Chemicon, #AB1778, 1:500), VIP (Rabbit, Immunostar, #20077, 1:400), NPY (Rabbit, Biogenesis, #6730-0004, 1:400), SYN1 (Rabbit, Abcam, #ab64581, 1:400), CSDE1 (Rabbit, Abcam, #ab201688, 1:200), GSK3β (Rabbit, Abcam, #ab32391, 1:200), PEG12 (Goat, RayBiotech, #NP_038816.1, 1:200), DPYSL2 (Rabbit, Proteintech, #14686, 1:300), DBI (Rabbit, Aviva Systems Biology, ARP33135_P050, 1:200), APOE (Mouse, Abcam, #ab1906, 1:200), β-CATENIN (Mouse, BD Transduction Labs, #610154, 1:300), TCF4 (Rabbit, CST, #2565S, 1:300), HMGA2 (Rabbit, Abcam, #ab97276, 1:300), IGF2BP2 (Rabbit, Abcam, #ab124930, 1:300), SPARC (Goat, R&D Systems, #AF942, 1:400), CRYAB (Mouse, Abcam, #ab13496, 1:250), FCGRT (Rabbit, Thermo Fisher Scientific, #PA5-42871, 1:300). Secondary antibodies were used at a dilution of 1:500 in the block buffer and included donkey anti-rat/rabbit/mouse/goat conjugated with AlexaFluors 488, 568, Cy5 or DyLight647 as required (Jackson ImmunoResearch or Thermo Fisher Scientific).

### Microscopy, image analysis and quantifications

Stained sections and/or whole mount guts were imaged with a X40 (1.75 NA) and/or X20 (1.0 NA) objectives with the upright Leica single/multiphoton TCS-SP5 confocal microscope, supported by the LAS AF software. Images of cell culture experiments were acquired using the upright Olympus Confocal Laser Scanning microscope FV3000 supported by the FV31S-SW software (Olympus) using standard excitation and emission lasers for visualizing various fluorophores (405, 488, 568, 647). Live-cell imaging was performed with the inverted Leica DMI 4000B microscope, fitted with Leica EL6000 light source and supported by the Micromanager (2.0) software^81^. In addition, the Hamamatsu ORCA-spark digital CMOS camera was used to visualise the reporter (tdT) with a standard filter.

Images were processed using ImageJ (Wayne Rasband, NIH) and/or Adobe Photoshop (Adobe systems).

In the deletion experiments of *Yap1* and *Wwtr1* genes, quantification was performed with ImageJ using the Cell Counter plugin to identify the proliferative (EdU+) and neurogenic (TuJ1+) potential of reporter (YFP+) labelled cells in each genotype, including the wild-type samples (four genotypes in total). Data have been obtained from a minimum of at least three independent experiments. For the early stages (E9.5-11.5), 30 sections of the embryo (n=6) from each genotype were prepared and every 5^th^ section was stained and analysed. Whole mount stainings were performed on at least n=3 animals of each genotype and imaged to perform quantification. For the late stages, at least n=4 guts were processed for whole mount immunostainings, followed by confocal imaging. Quantification of the proportion of EdU+ and TuJ1+ of the YFP+ cells was performed on the proximal, middle, distal and colonic segments of each gut.

In cell culture experiments, the field of view always included areas where enteric ‘ganglioids’ were visualised for quantification of cells.

### Patch clamp recordings of neurons in EGC cultures

Whole-cell patch clamp recordings were obtained at room temperature from the soma of visually identified glial-derived enteric neurons at DIV25-33. Cells were continuously perfused at 1.5mL/min with extracellular solution containing 140 mM NaCl, 4 mM KCl, 2 mM CaCl_2,_ mM MgCl_2,_ 11 mM glucose, 10 mM HEPES at pH 7.4. Patch electrodes (5±1 MΩ) were pulled from borosilicate glass and filled with the intracellular solution comprised of 135 mM K-gluconate, 10 mM KCl, 10 mM Na-phosphocreatine, 2 mM MgATP_2_, 0.3 mM Na_3_GTP, 10 mM HEPES at pH 7.25. Tetrodotoxin citrate (Abcam, ab120055) was added in some experiments to determine the nature of voltage-gated Na^+^ channels underlying cell activity.

Selected tomato-positive ‘neuronal’ cells were always part of a ‘ganglioid’ and identified with a larger cell size and diffuse tdT^+^ signal. Patch-clamped neurons were identified with the use of the Alexa Fluor 488 (20 μM) dye that was included in the pipetted solution. Signals were recorded with a Multi-clamp 700B amplifier and Digidata 4400a driven by PClamp 10.3 software (Molecular Devices). Data were sampled and digitized at 10 kHz. Passive properties were measured using a 100 ms, −5mV step from a holding potential of −60 mV. Active neuronal properties were probed in current clamp configuration using a series of 1s-long step current injections from −100 to 400 pA in 50 pA increments every 15s, whilst injecting sufficient current to maintain membrane potential at −60 mV. Data were analysed using Clampfit 10.7, Origin 2018b. Action potential properties were measured using the first action potential generated with a 100 pA current injection. Hyperpolarization-activated voltage sag was calculated as the difference between the peak hyperpolarization due to −100 pA current injection and the voltage at the end of the 1s step. After hyperpolarization (or AHP) amplitude was measured as the peak hyperpolarization following depolarizing current injections steps.

### Analysis of small intestine fluidigm scRNA-seq data

#### Alignment and quality control

Reads were aligned to the Ensembl GRCm38 genome using Tophat2^82^. Transcript counts, including counts for spliced and unspliced reads were obtained using the program Velocyto^83^.

Cells expressing more than 2000 genes and less than 10 percent mitochondrial genes were retained for further analysis. Contaminating epithelial cells (n = 9) were filtered from the data based on Cdh1 counts (>= 10 counts). Likewise, mesenchymal cells (n = 39) were filtered based on the expression of the marker Meis2 (>= 10 counts) and macrophages (n = 6) were filtered using expression of the marker Cd14 (>= 10 counts). After filtering 904 cells were retained, 143 cells (E13.5), 20 cells (E12.5 → E15.5), 129 cells (E17.5), 432 cells (P1), 11 cells (P26) and 169 cells (P61).

Genes detected in fewer than 3 cells or with total counts < 10 were omitted from further analysis (23299 genes were retained). Counts were normalised to counts per million (CPM) and log transformed (after the addition of a pseudocount of 1). Cell cycle scoring and assignment was performed in SCANPY^84^ using cell cycle genes from Tirosh et al, 2015^85^.

Batch effects were visualised using dimensionality reduction techniques (UMAP). This showed good mixing between batches for each time point, thus demonstrating that batch effects did not obscure the biology (Extended Data Fig. 1A).

#### Calculation of gene modules

Normalised gene dispersions were calculated using SCANPY^84^. Gene module analysis was performed using the package Antler (https://github.com/juliendelile/Antler^23^. Here all genes with a normalised dispersion > 0 were used as input to the algorithm; this was done to remove genes with a lower normalised dispersion than expected, which are likely to contribute noise to the analysis. The correlation and consistency cut-offs were set to 0.3 as these cut-offs resulted in compact and homogeneous groupings as revealed by a heatmap (Extended Data Fig. 3). The minimum number of genes a gene should be correlated with and the minimum number of cells with a “positive” binarised level per gene module were both set to three. The resulting modules appear biologically meaningful with a number of modules containing genes with similar known functions (e.g. cell cycle markers, glial markers, neuronal markers). Additionally, markers associated with excitatory and inhibitory neurons segregate to different modules. Functional analysis (GO biological process) was performed for each module using g:Profiler^86^, accessed programmatically using the gprofiler python module.

##### Dimensionality reduction and clustering

The 2954 genes identified as taking part in gene modules were used as input for PCA analysis in SCANPY^84^. The rationale behind the selection of these genes was that these are most likely to have biological relevance in the developmental process; a similar approach was taken by Delile et al.^23^. The first 15 PCs were selected for dimensionality reduction and clustering using the Louvain algorithm (resolution 1.5). The resulting clustering was manually curated and those clusters with similar transcriptional profiles that separated purely based on the expression of cell-cycle associated markers were merged.

To obtain an estimate of proportions of uncommitted cells and cells undergoing neurogenesis at each time point, kmeans clustering (k=2) was performed over the embryonic and P1 data on the normalised expression of 4 neurogenic (*Tubb3*, *Elavl4*, *Ret*, *Phox2b*) and 4 progenitor (*Erbb3*, *Sox10*, *Fabp7*, *Plp1*) markers. This resulted in a cluster with higher expression of neurogenic markers that included committed neuronal precursors, and one with higher expression of progenitor markers that included both gliogenic precursors and uncommitted progenitors.

##### Analysis of potential Csde1 regulation

The gene Csde1, which features in the identified gene module 16, and is highly expressed by eEPs and eENPs, is known as a post transcriptional regulator. To investigate the hypothesis that Csde1 could have a key regulatory role within its gene module we obtained iCLIP-seq data, which identified direct RNA targets of human CSDE1 in melanoma cells, from Wurth et al.^42^. Additionally, we obtained RNA-seq data on genes differentially expression in hESCs after CSDE1 knockdown from Ju Lee et al.^40^.

To map human gene names from these datasets to murine gene names, we used mappings from the NCBI HomoloGene database (ncbi.nlm.nih.gov/homologene). To test whether gene modules were enriched in RNA targets of Csde1 we used an upper tail hypergeometric test, with arguments to the R *phyper* function as follows:

**x** = the number of Csde1 RNA targets in the gene module

**m** = the number of Csde1 RNA targets in our whole dataset

**n** = the total number of genes in our dataset that are not Csde1 RNA targets and can be mapped to human homologues

**k** = the number of genes in the gene module which can be mapped to human homologues

To test whether gene modules are biased towards genes which are up- or down- regulated upon Csde1 knockdown, we used a Fisher exact test with the following contingency table:

|  | Downregulated | Upregulated |
| --- | --- | --- |
| In gene module |  |  |
| Not in gene module |  |  |

To investigate whether gene modules were enriched in transcripts with Csde1 binding motifs we used a script available published previously^41^. This scans transcripts for the consensus Csde1 binding sites [A/G]5AAGUA[A/G] and [A/G]7AAC[A/G]2 determined by SELEX^87^. Genes names from our data set were converted to RefSeq transcript IDs using the R biomaRt package. We considered a gene to contain a Csde1 binding site motif if any of its mapped transcripts contained such a binding motif. To assess whether modules were enriched in Csde1 binding motifs we used permutation testing.

### Pseudotime inference and velocity calculation

Pseudotime was inferred using Slingshot^20^, setting the starting point to the “eEP” cluster. The difference between the G2M phase and S phase was regressed out of the data prior to pseudotime analysis, to maintain differences between cycling cells and non-cycling cells but to regress out differences between cycling cells. Genes from the identified gene modules were again used as input to the PCA and the top 15 PCs used as features for calculation of pseudotime. Clusters which had been defined in “Dimensionality and clustering” were used as input to Slingshot, with the exception that the small EGC2 cluster was merged with the EGC1 cluster.

Smoothed profiles for the expression of transcription factors over pseudotime for neurogenic trajectories from branch points onwards were obtained by fitting spline curves (Monocle 2.12.0, genSmoothCurves function with 3 degrees of freedom)^24, 25^. At this stage any transcription factors expressed in less than 10 cells within the trajectory were filtered out. Differentially expressed genes over pseudotime were identified using Monocle’s differentialGeneTest function with gene level distributions modelled as negative binomial distributions with fixed variance (option expressionFamily=negbinomial.size()). Only embryonic and P1 time points (stages with active neurogenesis) were used for this analysis. Stochastic RNA velocity was calculated using the programme scVelo^21^. For inference of dynamics, velocity was calculated using genes identified as taking part in gene modules. These were filtered for genes with at least 20 counts and 10 unspliced counts. 1222 of these were identified to have significant velocity and were used for further calculations.

### Analysis of trajectory geometry

We asked whether changes in gene expression over pseudotime determine a well-defined direction, consistent throughout the course of a trajectory, or whether there are points where cells change course. To answer this question, we analysed the geometry of each trajectory, taking an approach similar to that described by Shapiro et al.^22^. Specifically, we compared the geometries of the trajectories from “eEP” to “AN” and “eEP” to “EGC”. Our rationale here being that both trajectories occupy the same amount of real time, with the initial cells having been sampled at embryonic day 13.5 and the final cells having been sampled from adult animals.

Each trajectory can be represented by 𝑛 points along pseudotime, 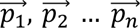 where each point is represented by coordinates along 𝑑 principal components, where 𝑑 is the number of dimensions considered. Unit vectors, representing the difference between the starting point 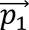 and each other point in the trajectory can then be calculated:

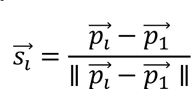

This results in 𝑛 − 1 unit vectors 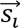 for each trajectory. If these are calculated over 3 dimensions (i.e. using the first 3 principal components), the resulting unit vectors can be visualised as points on a sphere. For a trajectory with no change in direction (i.e. all points in the trajectory fall on a straight line) each unit vector 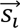 will be identical, and will project onto a single point on the sphere. In contrast, unit vectors calculated from a random trajectory, will disperse over the surface of the sphere. Thus the compactness of the area covered by a trajectory, projected onto the sphere, can be taken as a measure of how much the direction of a trajectory deviates throughout its course. Although easiest to visualise in 3 dimensions, this approach can be extended to an arbitrary number of dimensions.

To obtain points 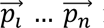representative of each trajectory, we employed a sampling strategy. First, we normalised pseudotime for each trajectory to range from 0 to 100. We then sampled a single cell from each sliding window of 10 pseudotime units. For each trajectory we repeated the sampling procedure 1000 times.

To assess the compactness of the area covered by the unit vectors 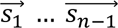 calculated for a trajectory, we found the unit vector 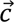, which minimised the mean spherical distance:

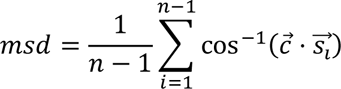

Minimisation was carried out using the “optim” function in R, using

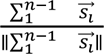

as a starting point for minimisation, and 1000 minimisation steps. The value for 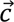 which minimises 𝑚𝑠𝑑 represents the direction of motion of the trajectory. We used the value 𝑚𝑠𝑑 for that 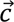 as a summary statistic to represent the compactness of the area covered by each trajectory.

We compared each empirical trajectory derived from the single cell transcriptomics data to a randomised trajectory. Here, the randomised trajectory is based on the empirical trajectory, however coordinates for each principal component are permuted (i.e. permutations are performed within each column of the matrix where rows represent cells and columns represent PCs).

Projections and summary statistics were calculated for randomised paths as for the empirical trajectories. The summary statistics for empirical (1000 sampled trajectories) and randomised (1000 randomised trajectories) data were then compared using Mann-Whitney U tests.

A representative centre 𝑐, which represents the direction of a developmental trajectory, was found by minimising 𝑚𝑠𝑑 for all centres calculated for the empirical trajectories. Genes were scored based on their association with this direction by calculating 𝐹 ⋅ 𝑐 where 𝐹 is the matrix of factor loadings for the 15 PCs used as input to Slingshot.

### Analysis of in vitro bulk data

‘Trim Galore!’ utility version 0.4.2 (https://www.bioinformatics.babraham.ac.uk/projects/trim_galore/) was used to remove sequencing adaptors and to quality trim individual reads with the q-parameter set to 20. The sequencing reads were then aligned to the mouse genome and transcriptome (Ensembl GRCm38 release-86) using RSEM version 1.3.0^88^ in conjunction with the STAR aligner version 2.5.2^89^. Sequencing quality of individual samples was assessed using FASTQC version 0.11.5 (https://www.bioinformatics.babraham.ac.uk/projects/fastqc/) and RNA-SeQC version 1.1.8^90^. Differential gene expression was determined using the R Bioconductor package DESeq2 version 1.14.1^91^.

### Analysis of in vitro time course 10X scRNA-seq data

Reads were aligned to the Ensembl GRCm38 using 10x Genomics Cell Ranger 3.0.2. Data was analysed in SCANPY. Cells with >1000 features and <10% mitochondrial counts were retained for downstream analysis. Predicted doublets were filtered using Scrublet^92^. Contaminating neurons (n=35) were filtered using the expression of the marker Elavl4 (> 0 counts) and contaminating macrophages (n = 99) were filtered using expression of the marker Cd14 (> 0 counts) from DIV0 data. Genes detected in fewer than 3 cells or with total counts < 10 were omitted from further analysis (23178 genes were retained). Counts were normalised to counts per million (CPM) and log transformed (after the addition of a pseudocount of 1). Cell cycle scoring and assignment was performed in SCANPY^84^ using cell cycle genes^85^. Highly variable genes were detected in SCANPY and the top 4397 used as input to PCA. The first 20 PCs were selected for dimensionality reduction and clustering using the Louvain algorithm (resolution 0.6). Marker genes for each cluster were detected using a Wilcoxon test, comparing each cluster to the union of all other clusters. Gene scores were calculated for selected Antler modules using SCANPY’s score_genes() function, and were visualised in SCANPY.

### Analysis of small intestine E15.5 10X scRNA-seq data

Single cell 10X data was obtained from GEO (GSE149524) for E15.5 Wnt-1-Cre mice^16^. Data was analysed using Seurat^93^. Cells with >1000 and <6000 features, <40000 counts and <5% mitochondrial counts were retained for downstream analysis. Counts were normalised to counts per million (CPM) and log transformed (after the addition of a pseudocount of 1). PCA was calculated using top 2000 most variable features and downstream clustering and UMAP were performed using 15 PCs. Louvain clustering was performed at a resolution of 0.8. Differential expression between clusters directly after the branch point (clusters 4 and 5; see Extended Data Fig. 4b) was performed in Seurat V4^93^ using the MAST package^94^ and the formula ∼cluster + nUMI.

### Analysis of autonomic fluidigm scRNA-seq data

Single-cell sequencing data (fastq files) were obtained from GEO (SRA) under accession GSE129114 for mouse E9.5 *Wnt1^Cre^/R26R^Tomato^*trunk cells. Autonomic cells were selected based on lineage annotations we reported earlier^3^. As for the small intestine developmental data, reads were aligned to the Ensembl GRCm38 genome using Tophat2^82^. Transcript counts, including counts for spliced and unspliced reads were obtained using the program Velocyto^83^. Counts were normalised to counts per million (CPM) and log transformed (after the addition of a pseudocount of 1). ANCC data was merged with the small intestine developmental data and the 2954 genes identified during the analysis of small intestine data were again selected for dimensionality reduction using PCA. The resulting PCA plot shows cells localise in a time dependent manner along PC1, with ANCCs localising at lower and overlapping values to E13.5 cells along this PC, reflecting a developmental progression. All ANCC batches localise together, and partially intermingle with cells from the closest time point (E13.5), indicating minimal batch effects. Differential expression between ANCCs and EGCs (union of P61 EGC1 and EGC2 cells), and between ANCCs and eEPs was performed in Seurat V3 using the MAST package^94^ and the formula ∼cell_type + nFeature_RNA

### Analysis of neural crest fluidigm scRNA-seq data

Single-cell sequencing data (raw counts) were obtained from GEO under accession GSE129114 for mouse E9.5 *Wnt1^Cre^/R26R^Tomato^* trunk cells. Counts were normalised to counts per million (CPM) and log transformed (after the addition of a pseudocount of 1) using Seurat V4^93^. Cells were labelled with lineage annotations we reported previously^3^.

### Analysis of neural stem cell data

Differentially expressed genes (microarray data) for quiescent and activated murine neural stem cells from the adult ventricular-subventricular zone were obtained from Codega et al.^69^. Similarly, differentially expressed genes (bulk RNA-seq) between cycling and cell cycle-arrested neural stem cells (a model of quiescence) in culture were obtained from Martynoga et al.^70^. To test whether the gene modules identified in the developmental data are enriched in genes that are upregulated in quiescent/cell cycle-arrested neural stem cells or activated/cycling neural stem cells, we used a Fisher exact test with the following contingency table:

|  | Upregulated in quiescent/cell cycle-arrested NSCs | Upregulated in activated/cycling NSCs |
| --- | --- | --- |
| In gene module |  |  |
| Not in gene module |  |  |

A Seurat object containing published pre-analysed scRNA-seq data for hippocampal neural stem cells was obtained from Harris et al.^71^. This contains neural stem cells from mice from 1 month to 6-8 months of age and includes dormant, resting and proliferating neural stem cells. Gene scores were calculated for selected Antler modules using SCANPY’s score_genes function and were visualised in SCANPY^84^.

### Analysis of scATAC-seq data

Reads for EGCs and neural crest cells (E9.5) were aligned to the mm10 genome using 10x Genomics Cell Ranger ATAC 1.2.0. Fragment files (output from 10x Genomics Cell Ranger ATAC 1.2.0) for the fresh cortex from adult mouse brain (P50) data were obtained from 10x genomics (https://www.10xgenomics.com/resources/datasets?menu%5Bproducts.name%5D=Single%20Cell%20ATAC&page=1&configure%5Bfacets%5D%5B0%5D=chemistryVersionAndThroughput&configure%5Bfacets%5D%5B1%5D=pipeline.version&configure%5BhitsPerPage%5D=500).

Downstream analysis was performed using ArchR^95^. Initially each dataset was investigated separately. Cells predicted to be doublets; cells with <= 10000 fragments; and cells with a TSS enrichment less than 4 were filtered from further analysis from each dataset. Iterative Latent Semantic Indexing (LSI) was performed and the first 30 dimensions used as input to the UMAP algorithm.

Cell types for the cortical and neural crest data were determined by label transfer using the Seurat FindTransferAnchors() function^93^ through the ArchR interface^95^. For the cortical dataset we downloaded annotated adult mouse cortical scRNA-seq data from the Allen Institute for Brain Science from https://www.dropbox.com/s/kqsy9tvsklbu7c4/allen_brain.rds in the form of a Seurat object. Single cell neural crest data was processed as in the section “Analysis of neural crest data”. EGC1 and EGC2 clusters in the scATAC EGC data were similarly inferred using our C1 fuidigm scRNA-seq EGC data.

For further analysis, cells labelled as oligodendrocytes and astrocytes from the cortical dataset were selected, and those labelled as belonging to the autonomic lineage were selected from the neural crest dataset. Iterative Latent Semantic Indexing (LSI) was performed on the entire dataset and the first 30 dimensions used to integrate the selected cortical, neural crest and EGC data using the Harmony algorithm^96^ via the ArchR interface^95^. Clustering was performed using the Louvain algorithm (resolution 1). Two small outlying clusters were removed and iterative LSI, integration using harmony and dimensionality reduction using UMAP were performed on this filtered dataset.

Gene scores were inferred using ArchR^95^; these provide an indication of chromatin acessibility at the gene level. Peaks were called using the “addReproduciblePeakSet()” function in ArchR^97^ that calls MACS2^98^. Cells were grouped by cell type for peak calling (autonomic cells, EGCs, oligodendrocytes, astrocytes). To determine the presence/absence of peaks in promoter regions, peaks were called individually for each cell type, using a q-value cut-off of 0.05.

To test whether gene modules were enriched in genes with at least one peak in their promoter region we used an upper tail hypergeometric test, with arguments to the R phyper function as follows:

**x** = the number of genes in the gene module with at least 1 peak in their promoter region.

**m** = the number of genes in the whole dataset with at least 1 peak in their promoter region.

**n** = the number of genes in the whole dataset without peaks in their promoter region.

**k** = the number of genes in the gene module.

Pairwise differential gene accessibility (using gene scores) between cell types was calculated using a wilcoxon test in ArchR with “TSSEnrichment” and “log10(nFrags)” as a bias. Here the bias features are quantile normalised, and for each cell in the group to be compared, ArchR selects its closest neighbour in the other group as a background. Pairwise differential peak accessibility was calculated in the same way.

Motif enrichments within differentially accessible peaks (logFC >= 1, padj <= 0.01) were calculated using ArchR for the Homer motif set^97^ (http://homer.ucsd.edu/homer/motif/). Additionally, chromVAR deviations^99^, bias-corrected measures of how far the per cell accessibility of a motif deviates from the expected norm, were calculated for Homer motifs in ArchR.

### Statistical Analysis

Bioinformatics statistical analysis was performed using R and python. All other statistical analyses were performed using GraphPad Prism 9.0 software (GraphPad). Normality distribution was tested using the D’Agostino– Pearson omnibus test. Unpaired two-tailed *t*-tests (followed by Welch’s correction test for non-equal standard deviations) were used to compare two groups. When comparing more than two groups, one-way ANOVA followed by Dunnett’s multiple-comparison test and Kruskal–Wallis test followed by a Dunn’s multiple-comparison test were used for parametric and non-parametric datasets, respectively. *P* < 0.05 was considered to be statistically significant and all of the *P* values are denoted in the figures. The nature of the entity of *n* is defined as individual animals (Fig. 2f, Fig. 4i, Extended Data Fig. 8p) or field of view (Fig. 4h, Extended Data Fig. 8n) and all experimental replicates are biological. All error bars represent mean ± standard error of the mean (s.e.m.) unless stated otherwise.

### Reporting summary

Further information on research design is available in the Nature Research Reporting Summary linked to this paper.

### Data availability

The .fastq files, SCANPY objects, count matrices and associated meta data will be made publicly available at the GEO repository. Source data will be provided with this paper.

### Code availability

The code used for all scRNA-seq analysis will be made available on GitHub.

## Acknowledgements

We thank the Crick Science Technology Platforms (STPs) for expert support, in particular the Biological Research Facility (BRF), the Flow Cytometry STP, the Advanced Sequencing Facility and the Experimental Histopathology Laboratory; We are grateful to Barry Thomson and Hannah Vanyai for helping us to establish colonies of the *Yap1^fl^* and *Wwtr1^fl^* alleles in our experimental unit in BRF. We also thank all members of the Pachnis laboratory for insightful comments on the manuscript. This work was supported by the Francis Crick Institute which receives its core funding from Cancer Research UK (FC001128, FC001159), the UK Medical Research Council (FC001128, FC001159), and the Wellcome Trust (FC001128, FC001159). V.P. acknowledges additional funding from BBSRC (BB/L022974) and the Wellcome Trust (212300/Z/18/Z).

## Notes

### Competing Interest Statement

The authors have declared no competing interest.

